# A Second Generation Leishmanization Vaccine with a Markerless Attenuated *Leishmania major* Strain using CRISPR gene editing

**DOI:** 10.1101/2020.05.04.077115

**Authors:** Wen Wei Zhang, Subir Karmakar, Sreenivas Gannavaram, Ranadhir Dey, Patrick Lypaczewski, Nevien Ismail, Abid Siddiqui, Vahan Simonyan, Fabiano Oliveira, Iliano V. Coutinho-Abreu, Thiago DeSouza-Vieira, Claudio Meneses, James Oristian, Tiago D. Serfim, Abu Musa, Risa Nakamura, Noushin Saljoughian, Greta Volpedo, Monika Satoskar, Sanika Satoskar, Pradeep K Dagur, J Philip McCoy, Shaden Kamhawi, Jesus G. Valenzuela, Shinjiro Hamano, Abhay Satoskar, Greg Matlashewski, Hira L. Nakhasi

## Abstract

Leishmaniasis is a debilitating and often fatal neglected tropical disease caused by *Leishmania* protozoa transmitted by infected sand flies. Vaccination through leishmanization with live *Leishmania major* has been used successfully but is no longer practiced because it resulted in unacceptable skin lesions. A second generation leishmanization is described here using a CRISPR genome edited *L. major* strain (*LmCen*^*-/-*^). Notably, *LmCen*^*-/-*^ is the first genetically engineered gene deleted *Leishmania* strain that is antibiotic resistant marker free and does not have any off-target mutations. Mice immunized with *LmCen*^*-/-*^ had virtually no visible lesions following challenge with *L. major*-infected sand flies while non-immunized animals developed large and progressive lesions with a 2-log fold higher parasite burden. *LmCen*^*-/-*^ immunization showed protection and an immune response comparable to leishmanization. *LmCen*^*-/-*^ is safe since it was unable to cause disease even in immunocompromised mice, induces robust host protection against vector sand fly challenge and because it is marker free, can be advanced to human vaccine trials.

## Introduction

Leishmaniasis is a neglected disease caused by infection with protozoans of the genus *Leishmania* that is transmitted by infected sand flies^1^. Worldwide, an estimated 1 billion people are at risk of infection in tropical and subtropical countries where up to 1.7 million new cases in 98 countries occur each year^2,3^. The disease pathology ranges from localized skin ulcers (cutaneous leishmaniasis, CL) to fatal systemic disease (visceral leishmaniasis, VL), depending on the species of the infecting *Leishmania* parasite^1,4^. Treatment options for both VL and CL are limited and there is poor surveillance in the most highly endemic countries^1,5^. A prophylactic vaccine would be an effective intervention for protection against this disease, reducing transmission and supporting the elimination of leishmaniasis globally. Currently there are no available vaccines against any form of human leishmaniasis.

Unlike most parasitic infections, patients who recover from leishmaniasis naturally or following drug treatment develop immunity against reinfection indicating that the development of an effective vaccine should be feasible^6–8^. Furthermore, leishmanization, a process in which deliberate infections with a low dose of virulent *Leishmania major* provides greater than 90% protection against reinfection and has been used in several countries of the Middle East and the former Soviet Union^9–11^. Leishmanization is however no longer practiced because it is ethically unacceptable due to the resulting skin lesions that last for months at the site of inoculation. The overall strategy of this study is to develop the next generation leishmanization that is safer by providing a protective immune response against cutaneous leishmaniasis without causing skin lesions.

In case of leishmaniasis cell mediated immunity is critical, and particularly, CD4 T cells play a crucial role in the protection against CL^12^. Specifically, host defense involves Th1 response due to T-cells primed by antigen presenting cells producing IL-12^13^. Production of IL-12 by antigen presenting cells and IFNγ by T cells are crucial for controlling the parasite numbers^13^. In contrast, Th2 cytokines, mainly IL-4, IL-5 and IL-13, an anti-inflammatory cytokine, suppress host immunity and help parasite survival while minimizing the tissue damage due to unchecked inflammation^13,14^. The differential effects of Th1 and Th2 dichotomy in cutaneous leishmaniasis is extensively studied in murine models^15^.

Studies with several candidate vaccines against CL including leishmanization demonstrated that the establishment of predominant Th1 type of immune response correlated with protection ^16–18^. In murine leishmanization models, it is well established that IFN-γ producing CD4 Th1 cells are essential in mediating protective immunity against re-infection^19,20^. Multifunctional effector Th1 cells which also produce high IFN-γ play a crucial role in host protection^21^. Recently it has been shown in leishmanized mice that rapidly recruited short lived effector T cells producing IFN-γ conferred significant level of protection and could be used as a biomarker of host protection^22,23^. These studies collectively show that any effective vaccine should similarly maintain these antigen specific CD4 T cell populations long enough to induce a robust protection against reinfection.

Centrin is a calcium binding protein and essential in the duplication of centrosomes in eukaryotes including Leishmania^24,25^. Previously, we have shown that *centrin* gene-deficient *Leishmania donovani* parasites are viable in axenic promastigote culture but do not proliferate in infected macrophages and are highly efficacious as a live vaccine in animal models^26–31^. However, using live-attenuated *L. donovani* as a vaccine in humans is high-risk because of the potential for visceralization resulting in fatal visceral disease. Further, previously generated gene deleted *L. donovani* strains required the incorporation of antibiotic resistance marker genes. The presence of antibiotic resistance genes in any attenuated live vaccine renders the vaccine unacceptable by regulatory agencies for human vaccine trials.

To overcome these drawbacks, we used CRISPR-Cas genome editing recently established for *Leishmania*^32–34^ to generate an attenuated *L. major centrin* gene deletion mutant (*LmCen*^*-/-*^). This represents a major milestone because *LmCen*^*-/-*^ is the first gene deleted *Leishmania* parasite to be developed containing no antibiotic resistant selection genes, an essential prerequisite for approval by regulatory agencies and advancement to human trials. *L. major* was used because this species is safer than *L. donovani* since *L. major* remains in the skin at the site of infection and does not cause visceral disease^1,4^. As demonstrated within, vaccination with *LmCen*^*-/-*^ is safe, immunogenic and protective against sand fly transmitted *L. major* infection, that mimics natural infection in highly relevant cutaneous leishmaniasis animal models meeting efficacy and ethical standards for advancement to human clinical studies.

## Results

### Generation and selection of centrin deficient L. major (LmCen^-/-^) by CRISPR-Cas

CRISPR-Cas genome editing has recently been developed to delete *Leishmania* genes with or without integration of antibiotic selection markers into the genome^32–34^. The experimental approach used to delete the *centrin* gene (Gene ID: LmjF.22.1410) from *L. major* is detailed in Figure 1. Two guide sequences targeted to the 5’ and 3’ flanking sequences of the *centrin* gene were designed and cloned into the *Leishmania* CRISPR vector pLdCNa&b (18, Figure 1A) and transfected into *L. major* (Friedlin V9) promastigotes. To delete the *centrin* gene sequence precisely at the locations determined by the 2 guide RNA sequences flanking the *centrin* gene without using marker gene replacement, a 50-nucleotide oligonucleotide donor DNA sequence was transfected into the promastigotes containing the CRISPR expression vector pLdCN as previously described^33^. The donor DNA consisted of 25 nucleotides 5’ from the upstream gRNAa cleavage site and 25 nucleotides 3’ from the downstream gRNAb cleavage site (Figure 1B). The exact targeted sequences flanking the *centrin* gene and diagnostic PCR primers are shown in Supplementary Figure 1A.

**Figure 1:**
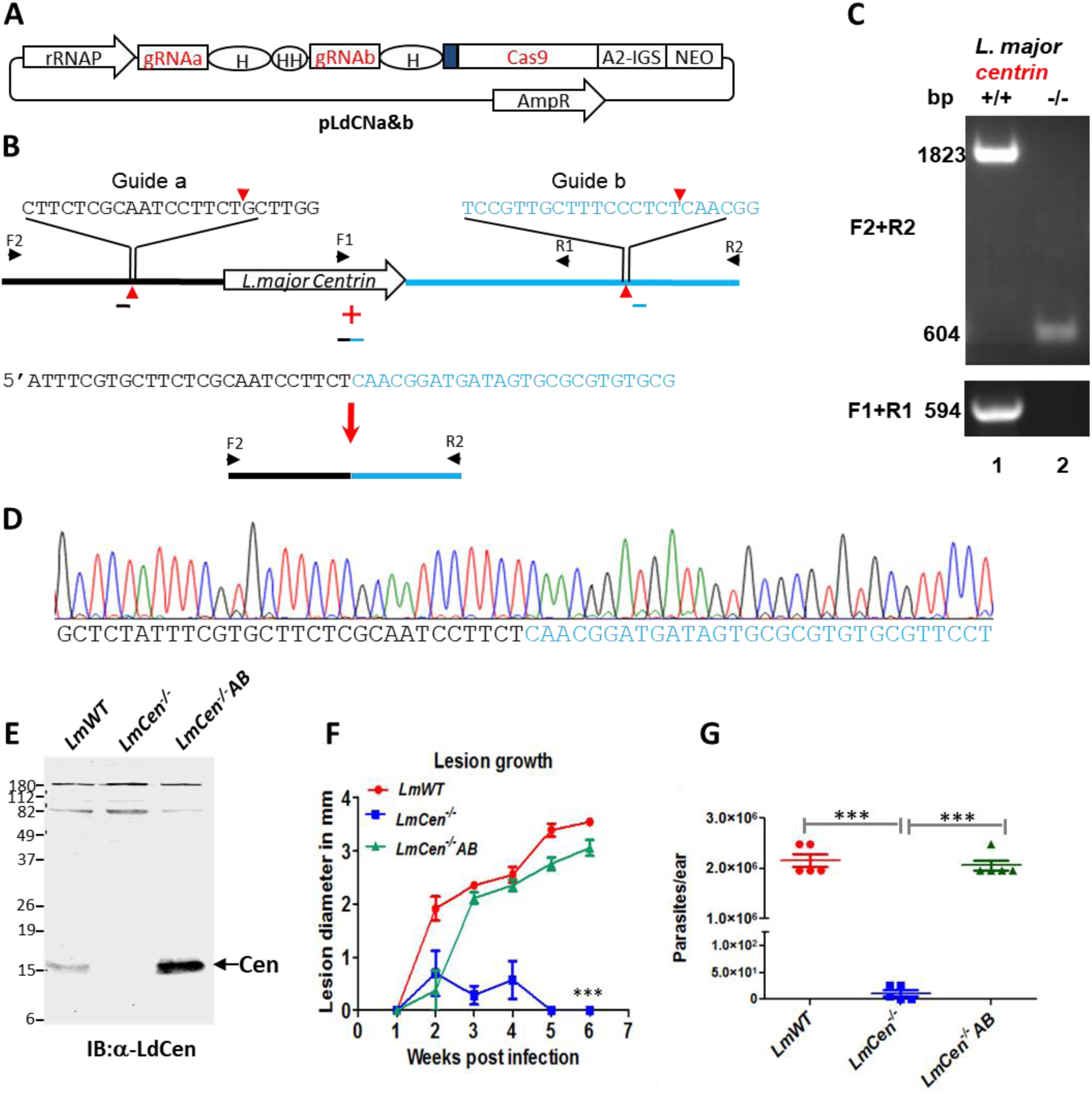
Generation of marker free *LmCen*^*-/-*^ parasite. Strategy for the generation of *centrin*-deficient *L. major* using CRISPR-Cas9. **A**. The pLdCN vector used to express Cas9 and gRNAa and gRNAb in *Leishmania*. A2-IGS, *L. donovani* A2 gene intergenic sequence; rRNAP, *L. donovani* ribosomal RNA promoter; H, Hepatitis delta virus ribozyme; HH, Hammerhead ribozyme. **B**. Schematic of gene deletion strategy showing gRNAa and gRNAb targeting sites in the *L. major centrin* gene locus and the expected gene deletion sequence after transfection of the cells with a 50 nucleotide oligonucleotide donor (18). The primers F1-R1 and F2-R2 used to detect this deletion are indicated. **C**. PCR analysis with primers F1-R1 and F2-R2 revealing loss of the *centrin* gene. Lane 1, Wildtype *L. major*; lane 2, *L. major centrin* null mutant. **D**. Sequence analysis confirming the flanking DNA breaks joined together by the transfected 50 nucleotide oligonucleotide donor. See the supplementary information for the detailed sequence. **E**. An immunoblot with an α-LdCentrin antibody showing the re-expression of Centrin in *LmCen*^*-/-*^ parasites transfected with a pKSNeo-LmCEN plasmid (*LmCen*^*-/-*^-AB, Addback). **F**. *LmCen*^*-/-*^ was unable to induce ear cutaneous lesions in C57BL/6 mice compared to wildtype *L. major* or the centrin add-back parasites of *LmCen*^*-/-*^ showing restored virulence (green line). C57BL/6 mice (n=5 per group) were infected intradermally (1 × 10^6^) with *LmWT, LmCen*^*-/-*^ or *LmCen*^*-/-*^*AB* parasites and the ear lesion development was monitored weekly. **G**. Parasite load in the infected ears of the mice. Parasite burden was determined by limiting dilution assay. Statistical analysis was performed by unpaired two-tailed t-test (***p<0.0001).

*L. donovani centrin* null promastigotes proliferate slower than wildtype promastigotes^35^. Since *centrin*-null promastigotes were selection marker free, this slower proliferation phenotype was used to identify *centrin* null *L. major* promastigotes. The CRISPR-genome edited *L. major* promastigotes were subjected to single cell cloning in 96 well plates; the relatively slow growing clones were identified, expanded and subjected to PCR analysis with the primers flanking the *centrin* gene as shown in Figure 1B. An example of a PCR analysis of a slow growing clone with the loss of the *centrin* gene is shown in Figure 1C. Sequence analysis of the 604 bp PCR product shown in Figure 1C confirmed the *centrin* gene containing sequence was precisely deleted at the predicted gRNA target sites and the chromosome fused through the donor sequence as intended (Figure 1D). The gRNA / Cas9 expressing pLdCN plasmid was subsequently removed from the *L. major centrin* null mutant (*LmCen*^*-/-*^) by single cell cloning and maintaining replica cultures in the presence and absence of G418 to identify clones sensitive to G418 that had lost the neomycin resistance gene present in the pLdCN CRISPR gene-editing plasmid. It was not possible amplify plasmid DNA from the G418 sensitive *LmCen*^*-/-*^ parasite providing further evidence for the loss of the pLdCN plasmid (Supplementary Figure 1B). As also shown in a Supplemental Figure 1C, the *LmCen*^*-/-*^ parasite retained the phenotype of slower proliferation than the WT *L. major*. This difference in proliferation enabled the identification and isolation of the slower growing *centrin* gene deleted clones by visual and microscopy inspection of the 96 well plate after one week in culture.

### LmCen^-/-^ promastigotes failed to produce lesions in infected mice

It was necessary to establish whether the *LmCen*^*-/-*^ had lost the ability to cause cutaneous infections and whether adding back the *centrin* gene through plasmid transfection (add-back, *LmCen*^-/-^*AB*) could restore cutaneous infection. The *centrin* gene was inserted into the *Leishmania* pKSNeo expression plasmid^36,37^, transfected into *LmCen*^*-/-*^ promastigotes and expression of the centrin protein was confirmed by Western blotting with an α-LdCen antibody that can recognize *L. major* centrin (Figure 1E). *LmCen*^*-/-*^ infection was investigated following intradermal injection of 1 × 10^6^ stationary phase promastigotes in the ear of C57BL/6 mice. As shown in Figure 1F, by 5-6 weeks, *LmCen*^*-/-*^ failed to produce swelling in the infected ear whereas wildtype *L. major (LmWT)* Friedlin V9 and the *LmCen*^*-/-*^ with the add-back *centrin* gene (*LmCen*^*-/-*^*AB)* did induce significant swelling. At 6 weeks following infection, the *LmCen*^*-/-*^ infected mice had few (<10) detectable parasites compared to both the *LmWT* and *LmCen*^*-/-*^*AB* infected mice that both had significantly more parasites (∼2 × 10^6^) (Figure 1G). These observations confirm that at 6 weeks post-infection, marker-free *LmCen*^*-/-*^ is unable to induce pathology at the site of injection in mice and that this was due to the deletion of the *centrin* gene.

We next examined *LmCen*^*-/-*^ survival in human macrophages *in vitro* since these are the obligate host cells for intracellular replication of *Leishmania* amastigotes (Supplementary Figure 1D). At 24 h post-infection, the number of parasites per macrophage was similar in the *LmCen*^*-/-*^ and *LmWT* infected cells. However, by 8 days, *LmCen*^*-/-*^ amastigotes were cleared from the macrophages, whereas *LmWT* parasites reached >10 parasites/macrophage. These results demonstrated that the *LmCen*^*-/-*^ promastigotes effectively infected human macrophages but subsequently were unable to proliferate intracellularly.

### LmCen^-/-^ contains no off-target gene deletions

Since the CRISPR generated *LmCen*^*-/-*^ strain was attenuated, it was necessary to establish the integrity of the genome by whole genome sequencing analysis to confirm the attenuation seen was solely due to the removal of the *centrin* gene. This analysis confirmed that the targeted ∼1kb genome region containing the 450 bp *centrin* gene (ID:LmjF.22.1410) was deleted from chromosome 22 and the remaining *centrin* gene homologs on chromosomes 7, 32, 34 and 36 remained intact in the genome (Figure 2A). Southern blot analysis confirmed the targeted *centrin* gene in *LmCen*^-/-^ was deleted and not translocated to another region of the genome (Figure 2B). Whole genome sequencing was performed to establish whether there were any off-target gene deletions in the edited genome. As shown in Figure 2C, the blue line is comprised of over 8,000 circles, each circle representing a single gene from chromosome 1 through 36 (left to right) whereas the red circles represent the members of the *centrin* gene family located on chromosomes 7, 22, 32, 34 and 36. There was virtually 100% coverage for all 8307 genes in the genome indicating the absence of partial or complete gene deletions, except the targeted *centrin* (LmjF.22.1410) that had a 0% coverage since it was deleted through CRISPR gene editing. A handful of genes (open blue circles) with less than 100% coverage are tandem repeat genes for which the coverage calculation software misaligned some reads, these genes were manually inspected and were found to be intact. Compared to the *L. major* Friedlin reference genome, there were no indels and no new SNPs (21 genes contained SNPs that were all previously identified in resequencing of the *L. major* Friedlin or LV39 strains). Collectively, these analyses demonstrate that the *LmCen*^*-/-*^ genome is intact and has no off-target gene mutations.

**Figure 2:**
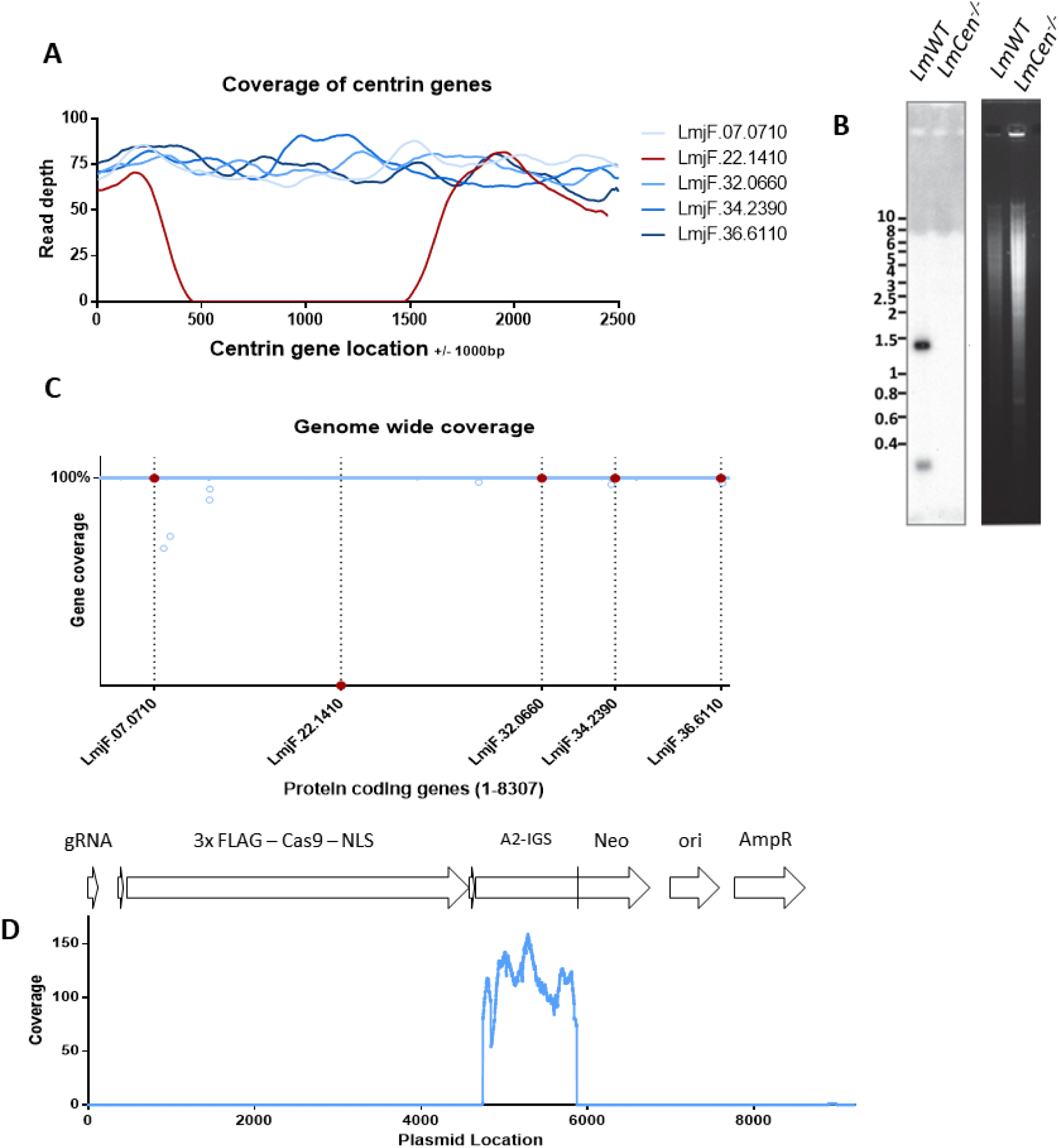
Whole genome analysis of the attenuated *LmCen*^*-/-*^ *L. major*. **A**. Sequence coverage across each of the *centrin* gene family members in the *LmCen*^*-/-*^ *L. major*. Note that only the targeted LmjF.22.1410 *centrin* gene has no sequence reads resulting from CRISPR gene editing. **B**. Southern blot analysis revealing the absence of the LmjF.22.1410 *centrin* gene in the genome of *LmCen*^*-/-*^ parasite compared to wildtype *L. major, LmWT*. **C**. Percent sequence coverage (Y-axis) for all protein coding genes from chromosome 1 to 36 (X-axis) by Illumina sequencing of the whole genome of the *LmCen*^*-/-*^ *L. major*. The blue line across the X axis is composed of 8307 dots where each dot represents a gene starting from chromosome 1 (left) to chromosome 36 (right) and is placed according to the portion of the open reading frame supported by sequencing reads. Open blue circles indicate genes where misalignments of sequencing Illumina reads occurred for some multicopy genes, although these genes were verified to be intact. Red circles and line markers correspond to the 5 *centrin* genes across the genome in chromosomes 7, 22, 32, 34 and 36. Only the targeted *centrin* gene (LmjF.22.1410) has been deleted from the genome and therefore has 0% coverage. D. Coverage of the pLdCN CRISPR plasmid sequence generated from whole genome sequencing. No homologous plasmid sequences were detected in the *LmCen*^*-/-*^ genome except for the positions ∼5000 to ∼6000 correspondings to the *Leishmania donovani* A2 gene intergenic sequence (A2-IGS) that were incorporated into the pLdCN plasmid for expression of the *Neo*^*R*^ gene. Therefore, the A2-IGS genomic sequence reads can align to this portion of the plasmid although the pLdCN CRISPR plasmid is not present in *LmCen*^*-/-*^.

The genomic DNA sequence reads were also searched for the presence of pLdCN CRISPR plasmid DNA sequences to confirm the loss of this plasmid. As shown in Figure 2D, the only *LmCen*^*-/-*^ genomic DNA sequences in common with the pLdCN CRISPR plasmid was the A2 gene intergenic sequence (A2-IGS) that is part of a A2 pseudogene sequence present in the *L. major* genome. The A2-IGS sequence from *L. donovani* was incorporated into the pLdCN CRISPR plasmid for processing of the Neo^R^ gene transcript^32^. There were no other detectable plasmid sequences or antibiotic resistance genes in the genome of *LmCen*^*-/-*^. It is noteworthy that the *L. donovani* ribosomal RNA promoter (rRNAP) sequence in the pLdCN CRISPR plasmid is sufficiently divergent from the *L. major* rRNAP sequence that it was not identified in the MiSeq DNA sequences by the Maximal Exact Match (bwa-mem) sequence alignment algorithm used. Taken together, the results presented in Figure 2D and Supplementary Figure 1B in combination with the loss of G418 resistance demonstrate that the pLdCN CRISPR gene-editing plasmid is no longer present in *LmCen*^*-/-*^. This represents a significant milestone since *LmCen*^*-/-*^ is the first marker free gene deleted *Leishmania* strain to be generated in the laboratory.

### Immunization with live LmCen^-/-^ is safe and does not cause lesions in highly susceptible mice

As shown in Figure 1, *LmCen*^*-/-*^ was unable to induce ear cutaneous lesions in C57BL/6 mice due to the removal of the *centrin* gene. However, to assess the safety of *LmCen*^*-/-*^ as a potential live vaccine, it was necessary to investigate its attenuation in a more susceptible mouse strain (BALB/c) and in immune deficient mice. BALB/c mice injected subcutaneously in the footpad with 1 × 10^7^ stationary phase *LmCen*^*-/-*^ showed no footpad swelling over 20 weeks (Figure 3A), the study endpoint, and a significantly lower parasite burden (approximately 4 log fold reduction) as compared to BALB/c mice injected with *LmWT* (Figure 3A). In some animals *LmCen*^-/-^ parasites were completely cleared by the study end point. Likewise, STAT-1 KO immune deficient mice injected with 2 × 10^8^ *LmCen*^*-/-*^ stationary phase parasites showed no footpad swelling during 7 weeks following injection (Figure 3B) whereas footpad swelling started at 4 weeks after injection with *LmWT* (Figure 3B). The parasite burden at 7 weeks in STAT-1 KO mice injected with *LmWT* was significantly higher than the mice injected with *LmCen*^*-/-*^ attenuated parasites (approximately 6 log fold reduction, Fig 3B). In another test, IFN-γ KO mice showed severe footpad swelling accompanied by a drastic increase in the number of the parasites after injection with 1 × 10^7^ *LmWT*, while injection with the same dose of *LmCen*^*-/-*^ did not show any footpad swelling in 20 weeks and the parasites were cleared from the site of injection (Figure 3C). The recombination activating gene 2 deficient (Rag2 KO) mice, which lack conventional T cells and B cells, showed mild footpad swelling and a high parasite burden in the footpad after 15 weeks following injection with *LmWT* (Figure 3D). In contrast, injection with *LmCen*^*-/-*^ did not show any swelling (Figure 3D; Supplementary Figure 2C) and the parasites were cleared from the site of injection (Figure 3D) and the spleen and liver (Supplementary Figure 2D). These results demonstrate that *LmCen*^*-/-*^ is non-pathogenic even in highly immunocompromised mice.

**Figure 3:**
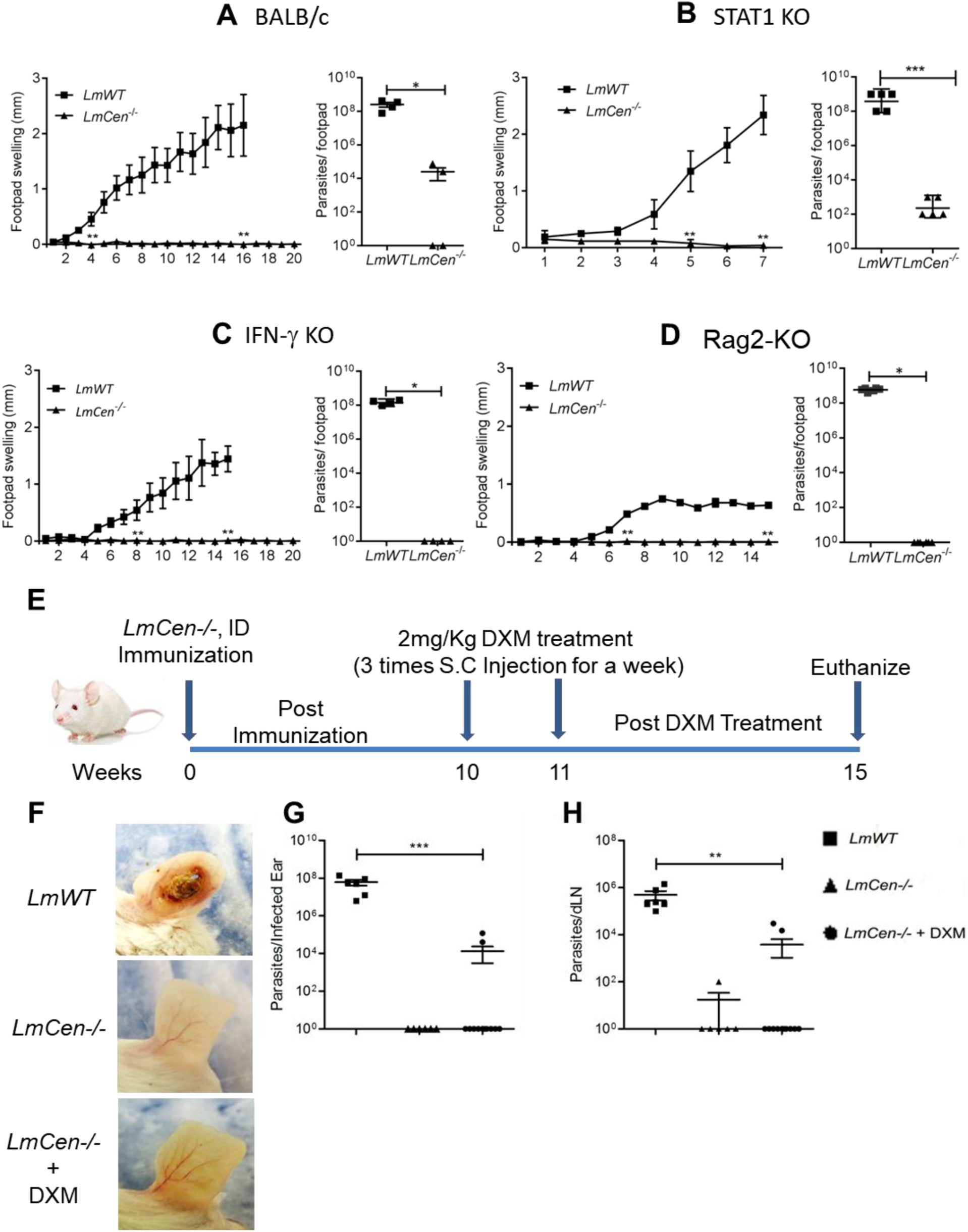
Safety and non-pathogenicity characteristics of *LmCen*^*-/-*^ parasites. **A**, BALB/c; **B**, STAT1 KO; **C**, IFN-γ KO and **D**, Rag2 KO mice were subcutaneously inoculated with indicated doses of *LmWT* or *LmCen*^*-/-*^ into the right hind footpad. **A, C, D**, BALB/c (n=4), IFN-γ KO (n=5) and Rag2 KO (n=6) mice were infected with 1 × 10^7^ of *LmWT* (Friedlin V9) or *LmCen*^*-/-*^, and (**B**) STAT1 KO (n=5) mice were infected with 2 × 10^8^ of *LmWT* (Friedlin V9) or *LmCen*^*-/-*^. Following infection, footpad swelling was measured weekly by digital caliper. Parasite burden in infected footpad was measured at 5 weeks after infection in BALB/c, at 7 weeks in STAT1 KO or at 15 weeks in IFN-γ KO and Rag2 KO. For the lesion development studies shown in A-D (left panels), asterisks represent the first time point at which significant differences were observed between the *LmWT* and *LmCen*^*-/-*^ groups. The differences in footpad swelling were statistically significant at all time points after the initial observation of the lesion. **E**, Schematic representation of the DXM treatment: BALB/c mice were divided into three groups. Groups-1 (n=6 mice/group) were infected with intradermal injection of 10^6^ total stationary phase *L. major* wildtype (*LmWT*) promastigotes in the ear dermis. Groups-2 (n=6 mice/group) and group-3 (n=12 mice/group) received an intradermal immunization of 1 × 10^6^ total stationary phase *centrin* deleted *L. major* (*LmCen*^*-/-*^) promastigotes in the ear dermis. Ten weeks post inoculation, only the third group was treated with 2 mg/kg DXM over one week by 3 subcutaneous injections. All three groups of animals were euthanized four weeks after DXM treatment (total 15 weeks post infection). **F**, Photographs of representative ear of *LmWT* infected (Group1) and *LmCen*^*-/-*^ (Group2) *(LmCen*^-/-^+DXM) (group3) mice. Compared to *LmWT* group which develops severe pathology in the ear, *LmCen*^*-/-*^ (±DXM) immunized mice display no ear pathology. **G** and **H**, Scatter dot plot of parasite load in infected ear (**G)** and draining lymph node (dLN) (**H**) of each *LmWT* and *LmCen*^*-/-*^ (±DXM) immunized mice. Parasite burden was determined by limiting dilution assay. Results represent data pooled from two independent experiments with (n=3-6 mice per group each time). Statistical analysis was performed by unpaired two-tailed t-test (**p < 0.004, ***p<0.0006).

To rule out the survival of any undetectable *LmCen*^*-/-*^ parasites beyond 7 weeks post-immunization, *LmCen*^*-/-*^ infected BALB/c mice were treated with 2 mg/kg dexamethasone (DXM), a known immune suppressor, three times for a week starting at 10 weeks post-infection (Figure 3E). All the groups were sacrificed at 4 weeks after the DXM treatment to determine parasite burdens. As shown in Figure 3F and Supplementary Figure 2A, *LmCen*^*-/-*^ infected mice with or without DXM treatment resulted in no lesions while *LmWT* infected but DXM-untreated mice developed open ulcerative lesions in the ear. Moreover, only 2 of 12 DXM-treated mice infected with *LmCen*^-/-^ showed parasites in the inoculated ear (Figure 3G) and draining lymph node (Figure 3H). In 1 of 6 untreated *LmCen*^*-/-*^-immunized animals, a low parasite number was detected in the draining lymph node (<100 parasites, Figure 3H), and none in the ear (Figure 3G). In *LmWT* infected mice, a significantly higher parasite load was observed in the ear and draining lymph node compared to *LmCen*^*-/-*^ -infected mice (± DXM) (Figure 3G, H), which correlated with ear lesion size (Figure 3F). Further, a PCR analysis using *L. major centrin* gene specific primers confirmed the absence of the *centrin* gene in the parasites isolated from DXM-treated mice (Supplementary Figure 2B lane 1, red arrow). Collectively, these results revealed that the centrin deleted live *LmCen*^*-/-*^ parasites are unable to revert or cause pathology and are safe for further study as a live vaccine.

### LmCen^-/-^ immunization induced protection against needle infection with wildtype L. major

To investigate the protective efficacy of *LmCen*^*-/-*^ against wildtype *L. major*, both resistant (C57BL/6) and susceptible (BALB/c) mice were immunized with a single intradermal (i.d.) injection with 1 × 10^6^ stationary phase *LmCen*^*-/-*^ in one ear. Seven weeks post-immunization, mice were challenged with 750 metacyclic wildtype *L. major* (WR 2885 strain) parasites in the contralateral ear via the i.d. route (Figure 4A for C57BL/6; Supplementary Figure 3A for BALB/c). Following challenge with wildtype *L. major*, lesion development was assessed up to 10 weeks for the C57BL/6 mice (Figure 4B, C). In the non-immunized-challenged group, mice developed a non-healing open ulcer that progressively increased in size (Figure 4C). No open ulcers were observed in the *LmCen*^*-/-*^-immunized-challenged group and only a moderate swelling that subsided from 5-9 weeks post-challenge was observed in 6 of 13 mice. Figure 4C depicts the ear pathology at 10 weeks post-challenge compared to a naïve unchallenged ear. Importantly, histopathological analysis revealed no clear difference between immunized-challenged and naïve mice ears, while non-immunized-challenged mice ears developed large lesions with open ulcers involving an influx of inflammatory cells (Figure 4C). The parasite load in the challenged ear and draining lymph node were also quantified at 10 weeks post-challenge revealing that the immunized group had a significantly lower parasite load (approximately a 4-log fold and a 3.2-log fold reduction, respectively) compared to the non-immunized group (Figure 4D, E). Similarly, highly susceptible BALB/c mice were protected following immunization with *LmCen*^*-/-*^ parasites (Supplementary Figure 3A-E). At 10 weeks post-challenge with wildtype *L. major* parasites, immunized BALB/c mice were protected as measured both by a reduced lesion size (Supplementary Figure 3B, C) and parasite burden (Supplementary Figure 3D, E) compared to non-immunized-challenged mice. A similar lack of non-healing open ulcer was observed in *LmCen*^-/-^ immunized BALB/c mice challenged with other wildtype strains of *L. major* such as *L. major* FV9 (Supplementary Figure 3F) and *L. major* LV39 (Supplementary Figure 3G).

**Figure 4:**
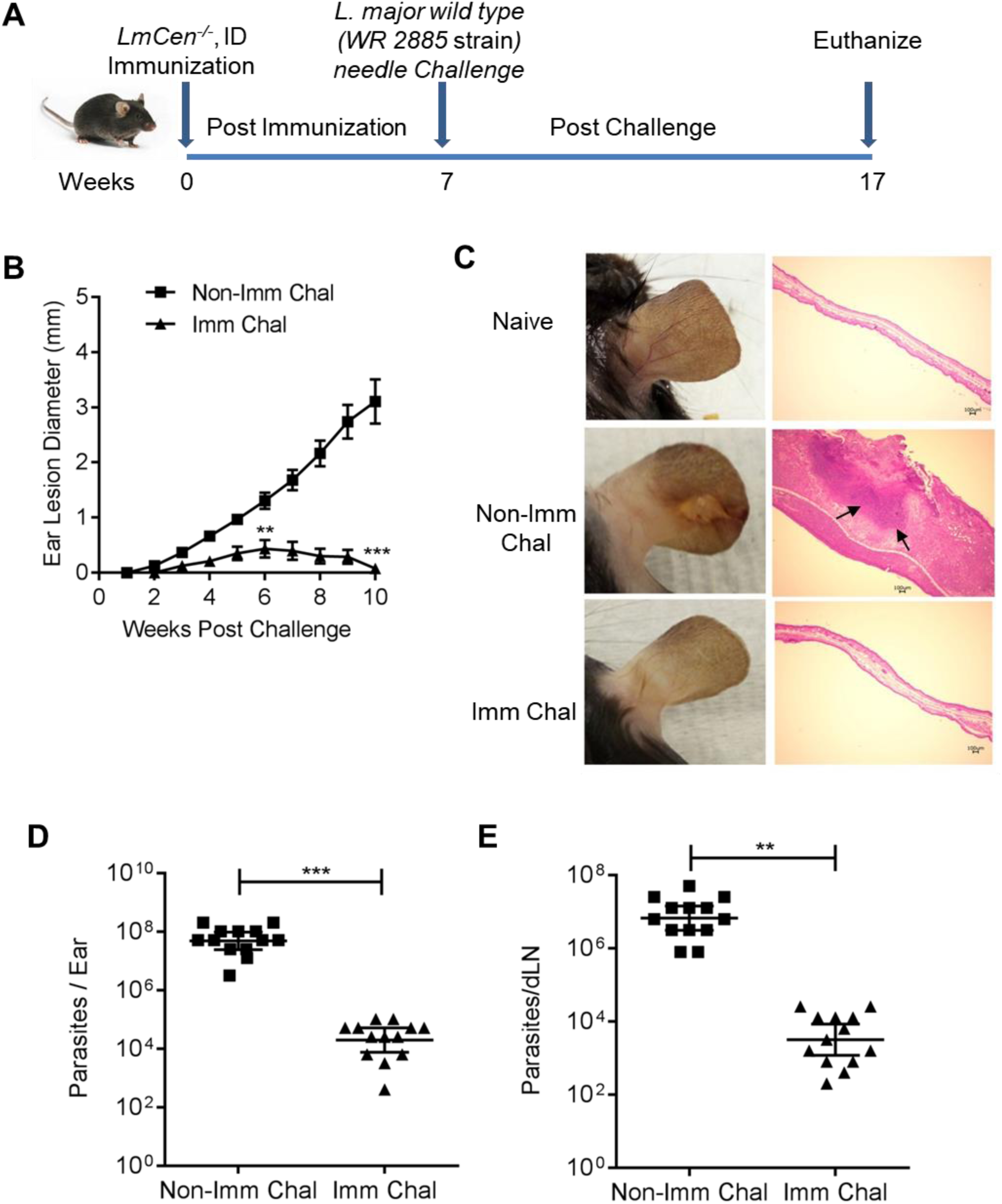
Protective efficacy of *LmCen*^*-/-*^ parasites against virulent *L. major* needle challenge in C57BL/6 mice. **A**. Schematic representation of the needle challenge procedure. Mice were immunized by intradermal injection in the left ear dermis with 1 × 10^6^ stationary phase *centrin* deleted *L. major* (*LmCen*^*-/-*^) promastigotes. Seven weeks post-immunization, both immunized and age matched naïve animals were challenged with 750 metacyclic *L. major* wildtype parasites in the right ear by intradermal injection. All the animals were euthanized after 10 weeks post challenge as shown in the figure. **B**. Ear lesion size was measured weekly for both *LmCen*^*-/-*^ immunized (Imm Chal) and non-immunized (Non-Imm Chal) mice after intradermal challenge with *LmWT* parasites. Results are mean ± SEM. For the lesion development studies shown in B, asterisks represent the first time point at which significant differences were observed between immunized (Imm Chal) and non-immunized (Non-Imm Chal) mice. The differences in ear lesion diameter were statistically significant at all time points after the initial observation of the lesion. **C**. Photographs (left panel) & histology (H&E stained, right panel) of representative challenged ear of *LmCen*^*-/-*^ immunized (Imm Chal) & non-immunized (Non-Imm Chal) mice after 10 weeks post challenge. Arrow indicates inflammatory cells recruited area. **D** and **E**, Scatter dot plot of parasite load of challenged ear (**D**) and draining lymph node (**E**) of each *LmCen*^*-/-*^ immunized (Imm Chal) & non-Immunized (Non-Imm chal) mice. Parasite burden was determined by limiting dilution assay. Results are mean ± SEM. Data are pooled from two independent experiments (n = 13 per group). Statistical analysis was performed by unpaired two-tailed t-test (**p < 0.001, ***p<0.0002).

### LmCen^-/-^ immunization induced protection against sand fly transmitted L. major infection

It is substantially more difficult and more relevant to demonstrate immunological protection against *L. major* infection initiated by a sandfly challenge than by a needle injection challenge^20,38^. Therefore, to determine the efficacy of *LmCen*^*-/-*^ immunization against sand fly transmitted cutaneous infection by *L. major*, C57BL/6 mice were immunized with a single i.d. injection of 1 × 10^6^ *LmCen*^*-/-*^ stationary phase parasites and mice were infected by exposure to bites of 10 *L. major*-infected sand flies in the contralateral ear 7 weeks post-immunization (Figure 5A). Disease progression was monitored for 10 weeks post-challenge by measuring lesion growth and assessing parasite burden in the ear and draining lymph node (Figure 5B-E). Notably, only 1/12 immunized-challenged mice developed a visible lesion, while 10/14 non-immunized-challenged mice developed progressive lesions in the ear that were significantly larger than the single lesion observed in immunized-challenged mice (Figure 5B, C). At 10 weeks post-challenge, there was a significant reduction of the parasite burden both in the ear and draining lymph node (approximately 2 log fold reduction in both) of immunized-challenged mice compared to non-immunized-challenged mice (Figure 5D, E). It is interesting to note that some of the draining lymph nodes in the immunized-challenged mice did not have any parasites (Figure 5E). These results demonstrate that immunization with *LmCen*^*-/-*^ mediates significant protection under natural conditions of infection i.e. parasite transmission by an infected sand fly.

**Figure 5:**
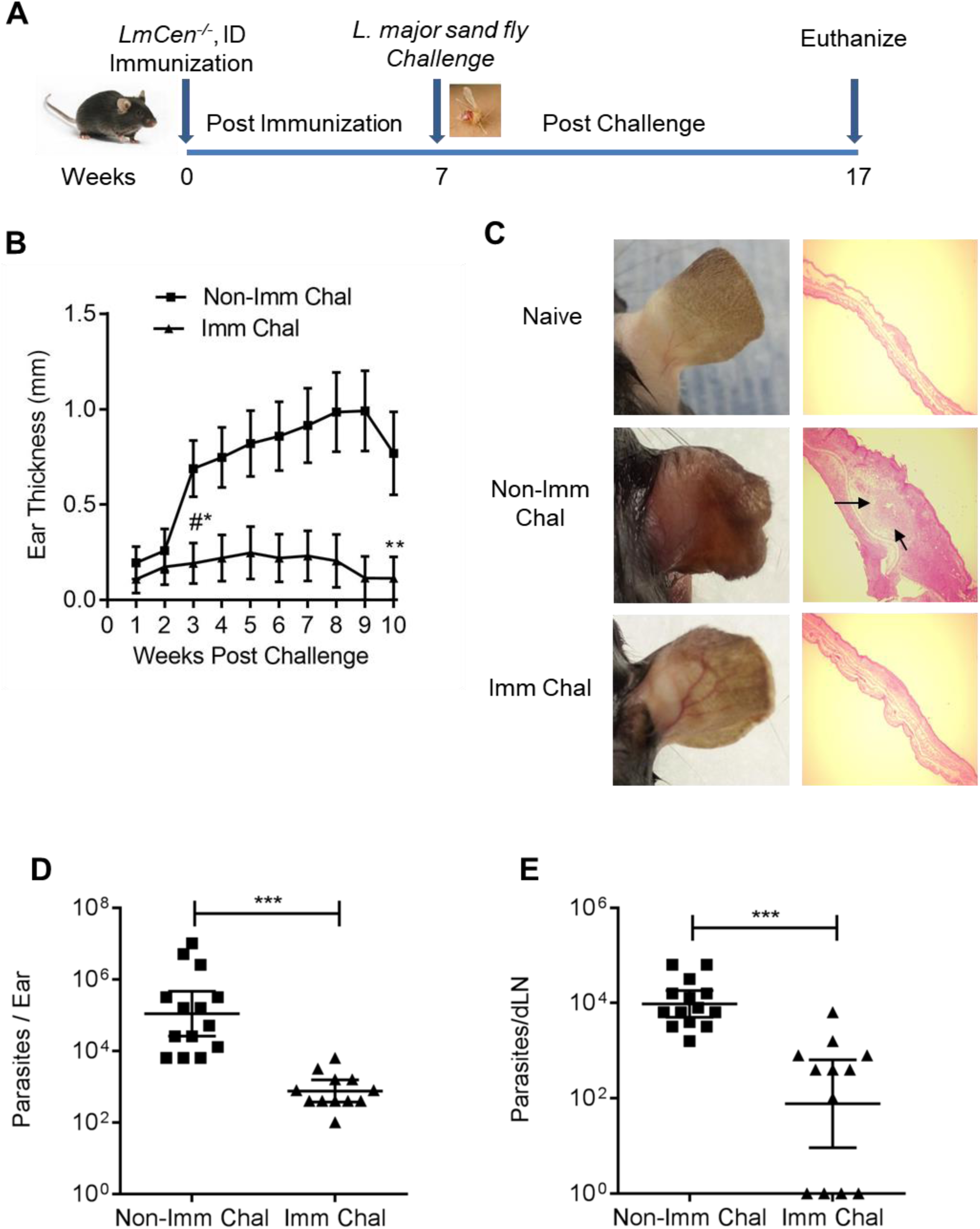
Protective efficacy of *LmCen*^*-/-*^ parasites against sand fly challenge in C57BL/6 mice. **A**. Schematic representation of the sand fly challenge procedure. Mice were immunized by intradermal injection in the left ear dermis with 1 × 10^6^ stationary phase *centrin* deleted *L. major* (*LmCen*^*-/-*^) promastigotes. Seven weeks post-immunization, both immunized and age matched naïve animals were challenged with ten *L. major* (WR 2855 strain) infected sand flies in the right ear. All the animals were euthanized after 10 weeks post challenge as shown in the Figure. **B**. Ear lesion thickness was measured weekly for both *LmCen*^*-/-*^ immunized (Imm Chal) and non-immunized (Non-Imm Chal) mice after sand fly transmission. only 1 mouse out of 12 *LmCen*^*-/-*^ immunized-challenged has developed severe lesion. For the lesion development studies shown in B, asterisks represent the first time point at which significant differences were observed between immunized (Imm Chal) and non-immunized (Non-Imm Chal) mice. The differences in ear lesion diameter were statistically significant at all time points after the initial observation of the lesion. **C**. Photographs (left panel) & histology (H&E staining right panel) of representative challenged ear of *LmCen*^*-/-*^ immunized (Imm Chal) & non-immunized (Non-Imm Chal) mice after 10 weeks post challenge. Arrow indicates inflammatory cells recruited area. Results are mean± SEM. **D** and **E**, Scatter dot plot of parasite load of challenged ear (**D**) and draining lymph node (**E**) of each *LmCen*^*-/-*^ immunized (Imm Chal) & non-Immunized (Non-Imm chal) mice. Parasite burden was determined by limiting dilution assay. Results are geometric means with 95% Cl of total 12-14 mice in each group. Data are pooled from two independent experiment. Statistical analysis was performed by Mann-Whitney two-tailed test (* p<0.05; ** p<0.01; ***p<0.0001).

### LmCen^-/-^ immunization or healed from primary infection with LmWT (leishmanization) induced comparable host protective immune response against L. major infection

Previously, in murine leishmanization models, it was shown that leishmanization induces host protective immunity against re-infection^19,20^. Having shown above that *LmCen*^*-/-*^ induces protection against both needle and the natural model of sand fly challenge, we first compared the immune response between *LmCen*^*-/-*^ immunized group (8 weeks of post-immunization) and a primary *LmWT* infection (healed group) at 12-weeks of post-primary infection (Figure 6A). The ears from the healed group showed lesions that later resolved. In the *LmCen*^*-/-*^ immunized group however, no lesion development was observed (Supplementary Figure 4A). Antigen experienced CD4^+^ T cells were first gated based on their surface expression of CD44 (Supplementary Figure 4B) and CD4^+^CD44^+^ cells were rearranged into different subpopulations based on their production of TNF-α, IFN-γ, and IL-2. The results showed that both *LmCen*^*-/-*^ immunization and healed groups of mice induced comparable single as well as multiple cytokines secreting CD4^+^CD44^+^T cells upon re-stimulation with *L. major* freeze-thaw antigen (*LmFTAg*) (Figure 6B & C). Naíve mice (not immunized with *LmCen*^*-/-*^ or infected with *LmWT)* did not show any detectable immune response after antigen stimulation (Figure 6B). Upon challenge with wildtype *L. major* parasites by needle injection, at 20hr post-infection, we observed a significant increase in the mRNA levels of IFN-γ in both healed and *LmCen*^*-/-*^ immunized ear tissues compared to nonimmunized mice (Figure 6D). From the same time point after challenge (20hr post-infection), we also analyzed the IFN-γ production from effector CD4 T cells by flow cytometry. Both healed and *LmCen*^*-/-*^ immunized mice induced a significantly higher percentage of IFN-γ^+^ effector T cells (CD4^+^CD44^+^T-bet^+^Ly6C^+^) compared to the non-immunized group (Figure 6E). Supplementary Figure 4C shows common gating strategies for early immune response (CD4^+^CD44^Hi^T-bet^+^Ly-6C^+^ IFN-γ^+^-T cells) in the ear of non-immunized, healed and *LmCen*^*-/-*^ immunized mice at 20hours post needle challenge with wildtype *L. major* parasites.

**Figure 6:**
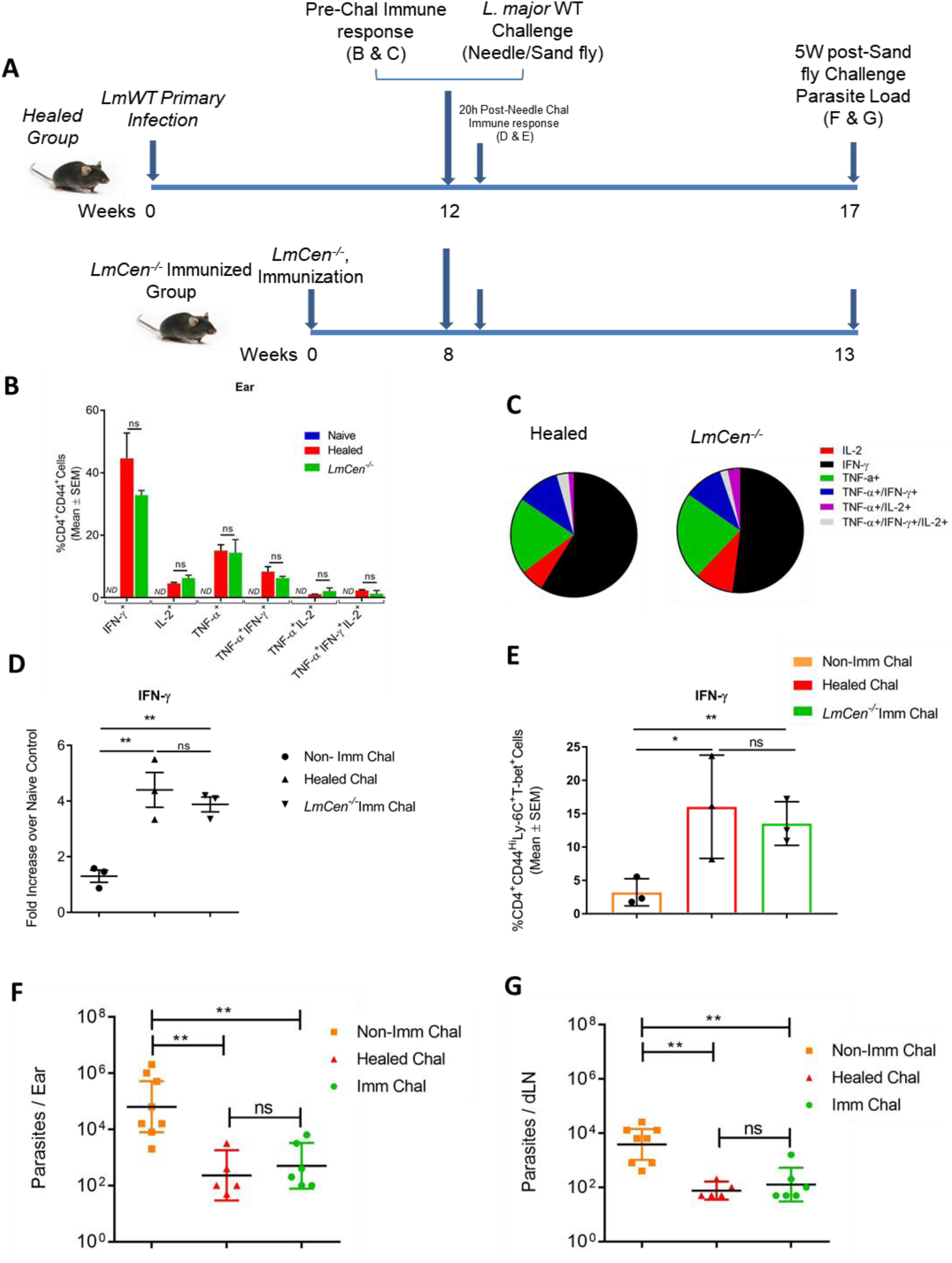
*LmCen*^*-/-*^ immunization or leishmanization with *LmWT* mediates comparable host protection against wildtype *L. major* infection. **A**. Schematic representation of the experimental approach used to comparative the protective immune response following leishmanization (healed) immunization with *LmCen*^*-/-*^. To determine comparative immune response between leishmanized and *LmCen*^*-/-*^ immunized mice (Fig. 6 B and C), C57BL/6 mice were either injected intradermally with 1 × 10^4^ metacyclic *LmWT* (Week-0) or 1 × 10^6^ total stationary phase *LmCen*^*-/-*^ parasites (Week-0) and a comparative immune response between healed from primary *LmWT* infection (Leishmanized) at week 12 and *LmCen*^*-/-*^ immunized mice at week 8 was determined. Naíve controls mice were not immunized with *LmCen*^*-/-*^ or not infected with *LmWT*. To determine the 20h post challenge immune response, healed, *LmCen*^*-/-*^ immunized as well as age matched naïve control mice were needle challenged with 1 × 10^5^ metacyclic *L. major* wildtype (*LmWT*) parasites in the contralateral ear (12 and 8 weeks respectively) (Fig. 6 D and E). To determine the protective response, healed, *LmCen*^*-/-*^ immunized and age matched naïve control mice (Fig. 6 F and G) were challenged with ten *L. major* infected sand flies in the right ear. All the animals were euthanized after 5 weeks post sand fly challenge (week 17 for leishmanized and weeks 13 for naïve control and *LmCen*^*-/-*^ immunized groups) and parasite load were determined. **B**. Multiparameter analysis for single, double or triple cytokine secreting CD3^+^CD4^+^CD44^+^ T cells after 20 hours of *in-vitro* re-stimulation with freeze-thaw *L. major* antigen (*LmFT*Ag) from pooled ears (2 ears) of naïve control, healed and *LmCen*^*-/-*^ immunized group of mice plus naive splenic APCs. Results (mean ± SEM) are representative of one independent experiment with 2-3 mice per group. Statistical analysis was performed by unpaired two-tailed t-test. **C**. Results were also represented in the Pie charts to show the cytokine profile of CD3^+^CD4^+^CD44^+^ T cells in response to *LmFT*Ag re-stimulation expressing any one cytokine (in red-IL-2, in black-IFN-γ and in green-TNF-α), any two cytokines (in violet – TNF-α^+^IL-2^+^ and in blue-TNF-α^+^IFN-γ^+^), all three cytokines (in gray-IL-2^+^TNF-α^+^IFN-γ^+^). The data presented are representative of single experiments. Mean and SEM of three mice in each group are shown. ns, p> 0.2. **D**. Ear IFN-γ expression were measured by RT-PCR analysis from healed, *LmCen*^*-/-*^ immunized and age matched naïve control mice following 20h post needle challenge with wildtype *L. major*-parasites. Results (mean ± SEM) are representative of two independent experiment with pooled ears (2 ears) samples (n=6 mice per group). ns, p> 0.48; **p> 0.009. **E**. Analysis of the early immune response following needle challenge with wildtype *L. major*-parasites. Twenty hours post-challenge, ear-derived cells were analyzed for IFN-γ producing CD3^+^CD4^+^CD44^hi^T-bet^+^Ly-6C^+^ T cells in response to 12-14 hours of *in-vitro* re-stimulation with freeze-thaw *L. major* antigen (*LmFT*Ag) plus naive splenic APCs. Results (mean ± SEM) are representative of two independent experiment with pooled ears (2ears) samples (n=6 mice per group). Statistical analysis was performed by unpaired two-tailed t-test (ns, p-0.31, *p> 0.02 and **p> 0.004). **F** and **G**. Five weeks of post-challenge with *L. major* WT infected sand fly, both ear (**F**) and draining lymph nodes (**G**) parasite load were determined by serial dilution. Results are geometric means with 95% Cl of total 5-8 mice in each group. Data are representative of one independent experiment. Statistical analysis was performed by non-parametric Mann-Whitney two-tailed test (ns, p-0.34; **p<0.004).

Healed and *LmCen*^*-/-*^ immunized groups were also challenged with *L. major* WT infected sand fly and the parasite loads were determined (Figure 6A). After five weeks of post-challenge, there was a similar significant reduction of parasite burden in the ear (2.4 log fold in healed group, and 2.1 log fold in *LmCen*^*-/-*^ immunized group) and draining lymph nodes (1.7 log fold in healed group, and 1.48 log fold in *LmCen*^*-/-*^ immunized group) compared to non-immunized group (Figure 6F and 6G). Both healed and *LmCen*^*-/-*^ immunized challenged mice did not develop any lesions whereas non-immunized challenged mice developed cutaneous lesions (Supplementary Figure 4D). Taken together, these results demonstrate that *LmCen*^*-/-*^ immunization is as effective as leishmanization (*LmWT* infection/healed) in generating a protective immune response and protecting against sand fly mediated infection with WT *L. major*.

## Discussion

Leishmanization with wildtype *L. major* has so far been the only successful human vaccine for leishmaniasis but it is ethically unacceptable because it causes skin lesions that last for months. This paper describes a second generation leishmanization live vaccination with an attenuated *L. major* strain (*LmCen*^*-/-*^) that does not cause lesions but retains the ability to provide immunological protection against experimental needle and sand fly transmitted *Leishmania* infection. As *LmCen*^*-/-*^ is marker gene free safe and efficacious, can be advanced to Phase I human clinical trials.

CRISPR-Cas genome editing was essential to generating this marker free strain because this technology can delete genes with high specificity and fidelity without selection with antibiotic resistant marker genes^32–34^. In place of antibiotic marker selection, the selection was based on a reduced proliferation rate of the *LmCen*^*-/-*^ mutant identified through single cell cloning, the first time such a selection has been performed in *Leishmania*. Whole genome sequence analysis confirmed that only the *centrin* gene on chromosome 22 (ID:LmjF.22.1410) was precisely deleted at the CRISPR guide RNA targeting sites and other *centrin* gene members on chromosomes 7, 32, 34 and 36 remained intact. There were no CRISPR-induced off-target gene deletions, indels or nonsynonymous SNPs introduced in the *LmCen*^*-/-*^ clone that was subjected to whole genome sequencing. By comparison, a previously engineered *L. donovani centrin* gene deleted parasites generated by homologous recombination with antibiotic resistant marker genes did contain off-target genomic deletions of up to 5000 base pairs in non-coding regions and in the coding regions of the folate transporter and gp63 genes^39^. Although gene-targeting specificity will depend largely on the selection of the guide RNA sequence, these observations suggest that CRISPR-Cas gene editing in *Leishmania* using a donor DNA fragment for repair as detailed in Figure 1 is more specific than traditional homologous recombination-based gene replacement with antibiotic resistance markers. In theory, it could have also been possible to generate a *centrin* gene deleted markerless *L. major* parasite using a different CRISPR approach involving the transfection of recombinant SaCas9 protein with *in vitro* –transcribed guide RNAs directed to upstream and downstream sequences flanking the *centrin* gene^40^. Although deletion of other *Leishmania* virulence genes may likewise generate attenuated strains, the *centrin* gene was targeted in this study because centrin gene deleted *L. donovani* parasites have been the most extensively validated parasites in previous experimental vaccine studies using various animal models^26–31^. It is noteworthy that in this study, *L. major* was used instead of the previous studies involving *centrin* deleted *L. donovani*^26–31^ as the focus of this study was cutaneous leishmaniasis.

Numerous experimental vaccines have been developed for *Leishmania*, though most of them have not been tested against natural sand fly transmitted infections. In studies when such vaccines were tested by needle challenge versus sand fly transmission of a virulent parasite, they were either partially protective or not protective against the latter^20,38,41,42^. In addition, sand fly mediated infection provides other components present in the saliva which play an important role in the pathogenesis of *Leishmania*^43–45^. The observations reported here demonstrated that markerless *LmCen-/-* immunization did induce protection against sand fly transmitted *L. major*. In this study, a major obstacle to using a live vaccine, the risk of disease development, was overcome by engineering a markerless second generation live attenuated parasite that can confer protection without associated pathology. Live attenuated *LmCen-/-* parasites elicited protective immunity in both susceptible (BALB/c) and resistant (C57BL/6) mice and against different strains of *L. major* (WR 2885, FV9 and LV39). Importantly *LmCen*^*-/-*^ parasites elicited protection against sand fly challenge that was deemed necessary but was neither performed or was not demonstrated in previous vaccination studies^20,38,41,42^. These observations using a cutaneous model of infection are consistent with our previous findings that immunization with *LdCen-/-* parasites were protective against visceral leishmaniasis in different animal models^26–28,30,46^.

In previous studies evaluating *Leishmania* vaccines, researchers have used mice with healed cutaneous lesions following a low dose of wildtype *L. major* infection as a gold standard animal model that mimics leishmanization in humans^23,38,47^. In this study, we have also compared *LmCen*^*-/-*^ parasite immunization induced immunity with wildtype *L. major* infected, healed mice (leishmanization). Our results demonstrated comparable immune responses in mice either healed from wildtype infection or immunized with *LmCen*^*-/-*^. It has been shown that chronic parasite infection maintains Ly6C^+^CD4^+^ effector T cells, and upon challenge with *LmWT* parasites these are essential for IFN-γ production that mediates protection^22^. Our results established that upon challenge with *LmWT* parasites, both *LmCen*^*-/-*^ immunized and healed mice generated a comparable percentage of CD4^+^Ly6C^+^IFN-γ^+^ effector T cells. In addition, protection may also be mediated by tissue resident memory T cells (Trm) that are called upon immediately after challenge as was shown in leishmanization mouse model^47^. Future studies with *LmCen*^*-/-*^ will address the role of Trm cell as well as other memory phenotype T cells in *LmCen*^*-/-*^ vaccine immunity. Moreover, upon *L. major* infected sand fly challenge, both groups are protected, and the levels of protection are comparable in terms of parasite burden. The residual parasite burden observed in both ear and lymph nodes in the *LmCen*^*-/-*^ may be important for maintaining long term protection as was reported in previous studies with leishmanized mice^22,48^. However, unlike leishmanization which involved inoculation of low dose of virulent parasites that caused lesions at the site of injection, immunization with *LmCen*^*-/-*^ parasites is safe as demonstrated by the absence of visible lesions in susceptible and immunodeficient animals post-immunization, in spite of persistence of a low number of *LmCen*^*-/-*^ parasites at the site of inoculation.

In conclusion, this study demonstrated that *LmCen*^*-/-*^ parasites are safe and can protect against a sand fly challenge with a wildtype *L. major* infection in relevant mouse models. Future studies are required to establish whether vaccination with *LmCen*^*-/-*^ is safe and protective in humans. The combination of old (leishmanization) and new (CRISPR gene editing) technologies can result in major advances in vaccine design that has the potential to protect millions of people from this major neglected disease.

## Materials and Methods

### Leishmania strain and culture medium

*L. major* Friedlin (FV9) and *L. major* LV39 used in this study were routinely passaged into the footpads of BALB/c mice. Amastigotes isolated from infected lesions were grown in M199 medium and promastigotes were cultured at 27°C in M199 medium (pH 7.4) supplemented with 10% heat-inactivated fetal bovine serum, 40 mM HEPES (pH 7.4), 0.1 mM adenine, 5 mg l^−1^hemin, 1 mg l^−1^ biotin, 1 mg l^−1^ biopterin, 50 U ml^−1^ penicillin and 50 µg ml^−1^ streptomycin. Cultures were passaged to fresh medium at a 40-fold dilution once a week. The growth curve of *L. major* promastigotes was obtained by inoculating the parasite at 1 × 10^6^ / ml into the 96 well plate (150 µl/well) in quadruplicate, the OD values were measured once a day for 4 days.

*L. major* WR 20885 strain was used to infect sand flies. This strain of parasites was isolated from a soldier deployed to Iraq and were grown at 27°C in Schneider’s medium supplemented with 10% heat-inactivated FCS, penicillin (100 U/ml), streptomycin (100 µg/ml), 2 mM l-glutamine. The WR2885 strain is shown to have superior colonization and transmissibility by sand flies to mice resulting in more severe pathology (larger lesion size and higher parasite loads)^38,49^.

### CRISPR Plasmid Construction

The pLdCNLm221410a&b plasmid vector was generated as follows: 1) A 276 bp PCR fragment containing gRNALm221410a, hepatitis delta virus and hammerhead ribozymes and gRNALm221410b guide coding sequences was amplified with primers Lm221410a and Ld221410b from the gRNA 241510+MT co-expression vector previously described^33,34^. The PCR product from step 1 was digested with Bbs I and inserted into the Bbs I digested pLdCN vector^33,34^ to generate the pLdCNLm221410a&b plasmid vector which was verified by sequencing analysis at the McGill University and Genome Quebec Innovation Center.

Guide RNA sequences and the oligonucleotide donor used in this study are listed below and their locations in the *centrin* locus are indicated in the Supplementary Figure 1.

gRNAa (Lm221410a): 5’ATCGAAGACCTTTGTCTTCTCGCAATCCTTCTGCTGTTTTAGAGCTAGAAATAGCA AG gRNAb (Lm221410b): 5’ATCGAAGACCCAAACTTGAGAGGGAAAGCAACGGACACCATGACGAGCTTACTC Oligo donor (Lm221410): 5’ATTTCGTGCTTCTCGCAATCCTTCTCAACGGATGATAGTGCG CGTGTGCG

### Selection of Centrin gene deleted clones and single cell cloning

*Leishmania* transfections were performed as previously described^37^. Briefly, 10 µg pLdCNLm221410a&b plasmid DNA was electroporated into 1 × 10^8^ early stationary phase *L. major* promastigotes. The transfected cells were then selected with G418 (100 µg/ml) for 2 weeks. Once the transfected *L. major* culture was established, the surviving promastigotes were subjected to three rounds of transfection with the oligonucleotide donor (Lm221410 oligo donor); 10 µl 100 µM single strand oligonucleotide donor was used per transfection, once every three days. After the third oligonucleotide donor transfection, the *Leishmania* promastigotes were counted and inoculated into 96 well plates at one promastigote per 100 µl medium per well. The growth of *Leishmania* cells in 96 well plates was monitored under microscope. After culture for three weeks in 96 well plates, parasites from the relatively slow growing clones were expanded in 24 well plates. The slow growing clones were selected since this represents the phenotype for loss of the *centrin* gene^35^. The genomic DNA extracted from the slow growth clones were subjected to PCR and DNA sequencing analysis to confirm deletion of the *centrin* gene.

To remove the pLdCNLm221410a&b plasmid from the *centrin* gene deleted *L. major* strain, individual clones were grown in duplicate plates where one plate contained media with G418 and the duplicate plate contained media without G418. Clones that had lost the plasmid were identified since they lost the ability to survive in the presence of G418.

### Genome sequence analysis of LmCen^-/-^

Complete genome sequencing of two clones from *LmCen*^*−/−*^ was determined by MiSeq genome sequencing reaction on an Illumina sequencing instrument at the sequencing core facility at the Center for Biologics Evaluation and Research. *LmCen*^*−/−*^ sequence reads were aligned against *Leishmania major* Friedlin strain reference genome (retrieved from www.tritrypdb.org) using the Burrows-Wheeler Aligner Maximal Exact Match algorithm (BWA-MEM)^50^. The alignments were converted to BED files using samtools and processed using the bedtools software package^51,52^. The bedtools coverage command was used with the “-d” option in conjunction with the genomic intervals containing the centrin genes to count the read depth at each position in the coverage of centrin genes shown in Figure 2A with a 200 bp window. The bedtools coverage command was used in conjunction with gene coordinates extracted from the gff genomic annotation file (retrieved from www.tritrypdb.org^53^) to compute the percent coverage of each gene as shown in Figure 2C. Genes with less than 100 percent coverage were manually inspected for a sharp drop-off in coverage (deletion) versus a gradual decline in close proximity to an inverse increase in coverage in a tandem gene (misalignment).

### Re-expression of centrin in LmCen^-/-^

The open reading frame encoding *centrin* gene was cloned into the SpeI sites of the *Leishmania* expression plasmid pKSNeo. *LmCen*^*-/-*^ parasites were transfected with the plasmid and recombinant parasites were selected using 50 µg/ml G418 to obtain *LmCen*^*-/-*^ parasites re-expressing centrin gene termed *LmCen*^*-/-*^ Addback (*LmCen*^*-/-*^AB).

### Southern hybridization

Total genomic DNA was isolated from promastigotes with the Wizard genomic DNA purification kit (Promega Biosciences). The DNA (5µg) was digested with restriction enzymes with BglI and the digestion products were separated on 1% agarose gels and transferred to positively charged nitrocellulose membranes. Southern blot analysis of the resolved DNA was performed as described previously using a ^32^p-labelled *L. major* centrin ORF nucleotide sequence as a probe^39^. The DNA fragments were ligated into pCR2.1-Topo vector and the nucleotide sequence of the probe was determined to ensure fidelity. The plasmid containing the correct probe was digested with EcoRI, gel purified and labeled with Random Prime it-II kit using ^32^p-dCTP (Agilent Technologies).

### Mice infection and immunization

Female 5-to 6-wk-old C57BL/6 and BALB/c mice were immunized and/or infected with 1 × 10^6^ total stationary phase *LmCen*^*-/-*^ or *L. major* wildtype (*LmWT*) parasites by intradermal injection in the left ear in 10 μl PBS. For challenge infections, age-matched naive and seven-week post immunized mice (both C57BL/6 and BALB/c) were challenged in the right ear with 750 metacyclic *L. major* (WR 2885 strain) wildtype promastigotes intradermally. The numbers of *L. major* (WR 2885) parasites in the infectious inoculum were determined by a titration analysis revealing that 750 metacyclic parasites cause reproducible pathology in BALB/c mice ear. For leishmanization, mice were infected with 1 × 10^4^ metacyclic promastigotes of *L. major* Friedlin (FV9) strain by intradermal needle injection in the ear. After 12 weeks of post-infection, healed mice were challenged on the contralateral ear with 1 × 10^5^ metacyclic *L. major* WR 2885 wildtype (*LmWT*) parasites by needle inoculation.

Lesion size was monitored up to 10 weeks post-challenge by measuring the diameter of the ear lesion using a direct reading Vernier caliper. Parasite burden in the challenged ear and draining lymph node (dLN) was estimated by limiting dilution analysis as previously described^37^. Briefly, two sheets of ear dermis were separated, deposited in DMEM containing 100 U/ml penicillin, 100 μg/ml streptomycin, and 0.2 mg/ml Liberase CI purified enzyme blend (Roche Diagnostics Corp.), and incubated for 1-2 h at 37°C. Digested tissue was processed in a tissue homogenizer (Medimachine; Becton Dickinson) and filtered through a 70 μm cell strainer (Falcon Products). Parasite titrations in the ear and dLN were performed by serial dilution (1:1 dilutions) of tissue homogenates in 96-well flat-bottom microtiter plates (Corning, Corning, NY) in M199 cell culture media in duplicate and incubated at 26°C without CO_2_ for 7–10 days. The greatest dilution yielding viable parasites was recorded and data are presented the mean parasite dilution ± SD. For histology, challenged ears were fixed, after 10 weeks of post WT parasite infection, in fixative solutions (10% buffered formalin phosphate solution) and paraffin-embedded sections were stained with hematoxylin and eosin (H&E) (Histoserv Inc.).

BALB/c mice were immunized subcutaneously in the footpad with 2 × 10^8^ *LmCen*^*-/-*^ parasites of the Friedlin strain or injected with PBS. After 6 weeks both groups were challenged with 10^4^ virulent metacyclics of LV39 *L. major* parasites intra-dermally in the ear. Ear lesions of vaccinated and non-vaccinated mice (PBS group) challenged with *L. major* LV39 metacyclic promastigotes were measured at least once a week from week 1 post challenge to week 10 post challenge.

BALB/c, IFN-γ KO and Rag2 KO mice were subcutaneously inoculated with 1 × 10^7^ of *LmWT* (Friedlin V9) or *LmCen*^*-/-*^ into the right hind footpad. Following infection, footpad swelling was measured weekly by digital caliper. Parasite burden in infected footpad was measured at 5 weeks after infection in BALB/c, at or at 15 weeks in IFN-γ KO and Rag2 KO mice. STAT-1 KO mice were injected subcutaneously in the footpad with 2 × 10^8^ *LmCen*^*-/-*^ parasites of the Friedlin strain or infected with 2 × 10^8^ *L. major* WT parasites of the Friedlin strain. Footpad swelling of both groups was measured at least once a week from week 1 after injection to week 7. After 7 weeks both groups were sacrificed, and parasite burden was determined. Footpad lesions was excised and then homogenized with a cell strainer in 3 ml of Schneider’s Drosophila medium (Gibco, US) supplemented with 20% heat-inactivated fetal calf serum and Penicillin-Streptomycin (0.1%).

### Sand fly infection and transmission of L. major to immunize mice

Female *Lutzomyia longipalpis* (Jacobina strain, reared at the Laboratory of Malaria and Vector Research, NIAID) sand flies were infected by artificial feeding through a chick skin membrane on a suspension of 5 × 10^6^ *L. major* (WR 2855 strain) procyclic promastigotes/ml of heparinized defibrinated blood containing penicillin and streptomycin. Flies with mature infections were used for transmission^54^. One day before transmission the sucrose diet was removed. Mice were anesthetized by intraperitoneal injection of 30 μl of ketamine/xylazine (100 mg/ml). Ointment was applied to the eyes to prevent corneal dryness. Ten infected flies were applied to right ears of both *LmCen*^*-/-*^ immunized and age-matched naïve C57BL/6 mice through a meshed surface of vials which were held in place by custom made clamps. The flies were allowed to feed on the exposed ear for a period of 2–3 h in the dark at 23°C and 50% humidity. Following exposure, the number of flies per vial with or without a blood meal was counted to determine the influence of feeding intensity on transmission frequency. Animals were sacrificed after 10 weeks of post sand fly exposure & organ parasite burden were determined by serial dilution as described above.

### Human macrophage infection

Human elutriated monocytes obtained from NIH blood bank from healthy US blood donors. Only monocytes that tested CMV negative were used in this study. Monocytes were re-suspended at 2 × 10^5^ cells/ml in RPMI medium containing 10% FBS and human macrophage colony-stimulating factor (20 ng/ml, ProSpec), plated in a volume of 0.5 ml in eight-chamber Lab-Tek tissue culture slides (Miles Laboratories) and incubated for 7 days for differentiation into macrophages. The differentiated macrophages were infected with stationary phase *LmWT* or *LmCen*^*-/-*^ promastigotes (10:1 parasite-to-macrophage ratio). After incubation for 6 h at 37°C in 5% CO_2_, the free extracellular parasites were removed by RPMI washes and the cultures were incubated in macrophage culture medium for an additional 24 h. The culture medium was removed, and macrophages infected with *LmWT* or *LmCen*^*-/-*^ were stained with Diff-Quik staining reagent. Percentages of infected macrophages were determined by counting a minimum of 100 macrophages per sample under the microscope. Results are shown as mean ± SEM for three independent counts for each infection on days 1-8.

### RT-PCR

Total RNA was extracted from the ears tissue using a Pure Link RNA Mini kit (Ambion). Total RNA (400ng) was reverse transcribed into cDNA using random hexamers with a high-capacity cDNA reverse transcription kit (Applied Biosytems). Gene expressions were determined using TaqMan Gene Expression Master Mix and premade TaqMan Gene Expression assays (Applied Biosystems) using a CFX96 Touch real-time system (Bio-Rad, CA) and the data were analyzed with CFX Manager software. The TaqMan Gene Expression Assay ID (Applied Biosystems) of IFN-γ (Mm01168134_m1) and GAPDH (Mm99999915_g1). Expression values were determined by the 2^−ΔΔCt^ method where samples were normalized to GAPDH expression and determined relative to naive sample.

### Measurement of cytokine expression from ear derived CD4^+^ T cell populations by flow cytometry

To determine the comparative immune response at pre- or 20 h post-*L. major* WT needle challenge, single-cell suspensions from ear of healed (leishmanized) and *LmCen*^*-/-*^ immunized mice were incubated with 1 × 10^6^ T-cell depleted (Miltenyi Biotech) naïve spleen cells (APCs), with 50 µg/ml freeze-thaw *L. major* antigen (*LmFT*Ag) in flat bottom 48-well plates at 37°C for 12-14 h. During last 4 h of culture, protein Transport Inhibitor (BD Golgiplug, BD Bio-Sciences) was added to the wells. Cells were then blocked at 4°C with rat α-mouse CD16/32 (5 µg/ml) from BD BioSciences for 20 min. For surface staining, cells were then stained with α-mouse CD3 AF-700 (BD BioSciences), α-mouse CD4 BV-650 (Biolegend) and α-mouse CD44 FITC (BD BioSciences) or α-mouse CD3 BV421 (BD BioSciences), α-mouse CD4 BV-650 (Biolegend), α-mouse Ly-6C APC-Cy7 (BD BioSciences) and α-mouse CD44 FITC (BD BioSciences) for 30 min (each with 1/300 dilution; 4°C). The cells were then stained with LIVE/DEAD fixable aqua (Invitrogen/Molecular Probes) to stain dead cells. Cells were washed with wash buffer and fixed with the Cytofix/Cytoperm Kit (BD Biosciences) for 20 min (room temperature). Intracellular staining was done with α-mouse IL-2 APC (BD BioSciences), α-mouse IFN-γ PE-Cy7 (Biolegend) and α-mouse TNF-α PerCP-Cy5.5 (Biolegend), for 30 min (each with 1:300 dilution; 4°C). In some experiments samples are treated with Foxp3 Fixation / Permeabilization Buffer (ebioscience) and then stained with α-mouse T-bet -BV786 (Biolegend) according to manufacturer’s instruction. Cells were acquired on Symphony (BD Biosciences, USA) analyzer equipped with 350, 405, 445, 488, 561, 638 and 785 nm LASER lines using DIVA software (v8). Data were analyzed with the FlowJo software version 9.9.6 (BD, San Jose CA). For analysis, first doublets were removed using width parameter; dead cells were excluded based on staining with the Live/Dead Aqua dye. Lymphocytes were identified according to their light-scattering properties. CD4^+^ T-cells were identified as CD3^+^ lymphocytes uniquely expressing CD4. Upon further gating intracellular cytokines were measured in CD44^hi^Ly-6C^+^T-bet^+^ cells. Fluorescence minus one control was used for proper gating of positive events for designated cytokines.

### Immunosuppression by dexamethasone injection

To determine the safety of Centrin deficient *LmCen*^*-/-*^ parasites in immune-suppressive condition, 4 to 6 weeks old BALB/c mice were divided into three groups. Group-1 (n=6) were infected with 1 × 10^6^ stationery phase *LmWT* parasites and Group-2 (n=6) and Group-3 (n=12) animals were immunized with 1 × 10^6^ stationery phase *LmCen*^*-/-*^ parasites in a 10μl volume of PBS through intradermal (into the ear dermis) routes. After 10 weeks of post infection, only Group-3 animals were treated with 2 mg/kg Dexamethasone sodium phosphate (Sigma Aldrich) in PBS by subcutaneous injection three times for one week. Four weeks after this treatment (total 15 weeks post infection); all the groups were sacrificed and evaluated for parasite burden by serial dilution as described above. Development of pathology & lesion size in the ear was assessed at 15 weeks post infection by measuring the diameter of the lesion.

Characterization of *centrin* deleted parasites isolated from *LmCen*^*-/-*^ plus DXM treated group was done by Polymerase chain reaction. Total Genomic DNA was isolated from the parasites recovered from *LmWT* and *LmCen*^*-/-*^ plus DXM treated group according to the manufacturer information (DNeasy Blood & Tissue Kit, Qiagen). PCR was performed with *L. major centrin* gene specific primer (For-5’-ATGGCTGCGCTGACGGATGAACAGATTCGC-3’; Rev-5’-CTTTCCACGCATCTGCAGCATCACGC-3’) which target the amplification of the 450-bp. A reaction mixture was prepared containing 10 × Buffer (Invitrogen), 0.2 mmol/l each deoxyribonucleotide (Invitrogen), 1 μmol/l each primer, 1.25 units of Taq polymerase (Invitrogen) and 200 ng of DNA samples in a final volume of 50 μl. The PCR conditions were as follows: denaturation at 94°C for 3 min, followed by 35 cycles of 94°C for 20 s, 58°C for 20 s and 68°C for 35 s with a final extension of 68°C for 5 min. The amplification reactions were analyzed by 1% agarose gel electrophoresis, followed by ethidium bromide staining and visualization under UV light. DNA from the reference plasmid (PCR 2.1 TOPO) containing *centrin* gene was used as a positive control.

### Statistical analysis

Statistical analysis of differences between means of groups was determined by unpaired two-tailed Student t test, using Graph Pad Prism 5.0 software. *, <0.05; ** < 0.005; and *** <0.0005 was considered significant.

### Ethical Statement

The animal protocol for this study has been approved by the Institutional Animal Care and Use Committee at the Center for Biologics Evaluation and Research, US FDA (ASP 1995#26). The animal protocol is in full accordance with “The guide for the care and use of animals as described in the US Public Health Service policy on Humane Care and Use of Laboratory Animals 2015”. All animal studies at Ohio State University were performed in accordance with NIH guidelines for the humane care and use of animals and were approved by OSU IACUC. Animal experimental procedures performed at the National Institute of Allergy and Infectious Diseases (NIAID) were reviewed by the NIAID Animal Care and Use Committee under animal protocol LMVR4E. The NIAID DIR Animal Care and Use Program complies with the Guide for the Care and Use of Laboratory Animals and with the NIH Office of Animal Care and Use and Animal Research Advisory Committee guidelines. Detailed NIH Animal Research Guidelines can be accessed at https://oma1.od.nih.gov/manualchapters/intramural/3040-2/. Animal experimental procedures performed at Nagasaki University were approved by the Institutional Animal Research Committee of Nagasaki University (No.1606211317 and 1505181227), the Nagasaki University Recombinant DNA Experiments Safety Committee (No. 1403041262 and 1407221278), and performed according to Japanese law for the Humane Treatment and Management of Animals.

## Author contributions

WWZ, SG, RD, SK, MS and SS designed and conducted experiments, analyzed data and helped write the manuscript. GM, HLN, AS, SH, SK, JGV designed experiments, analyzed data and wrote the manuscript. PL analyzed genome data and helped write the manuscript. NI, AS, VS, FO, IVC, TS, CM, AM, RN, NS, GV conducted experiments, analyzed data and reviewed the manuscript.

## Data availability statement

The data that support the findings of this study are available from the corresponding author upon reasonable request.

