## Supplementary Figures for "A Second Generation Leishmanization Vaccine with a Markerless Attenuated *Leishmania major* Strain using CRISPR gene editing"

### Supplementary Material

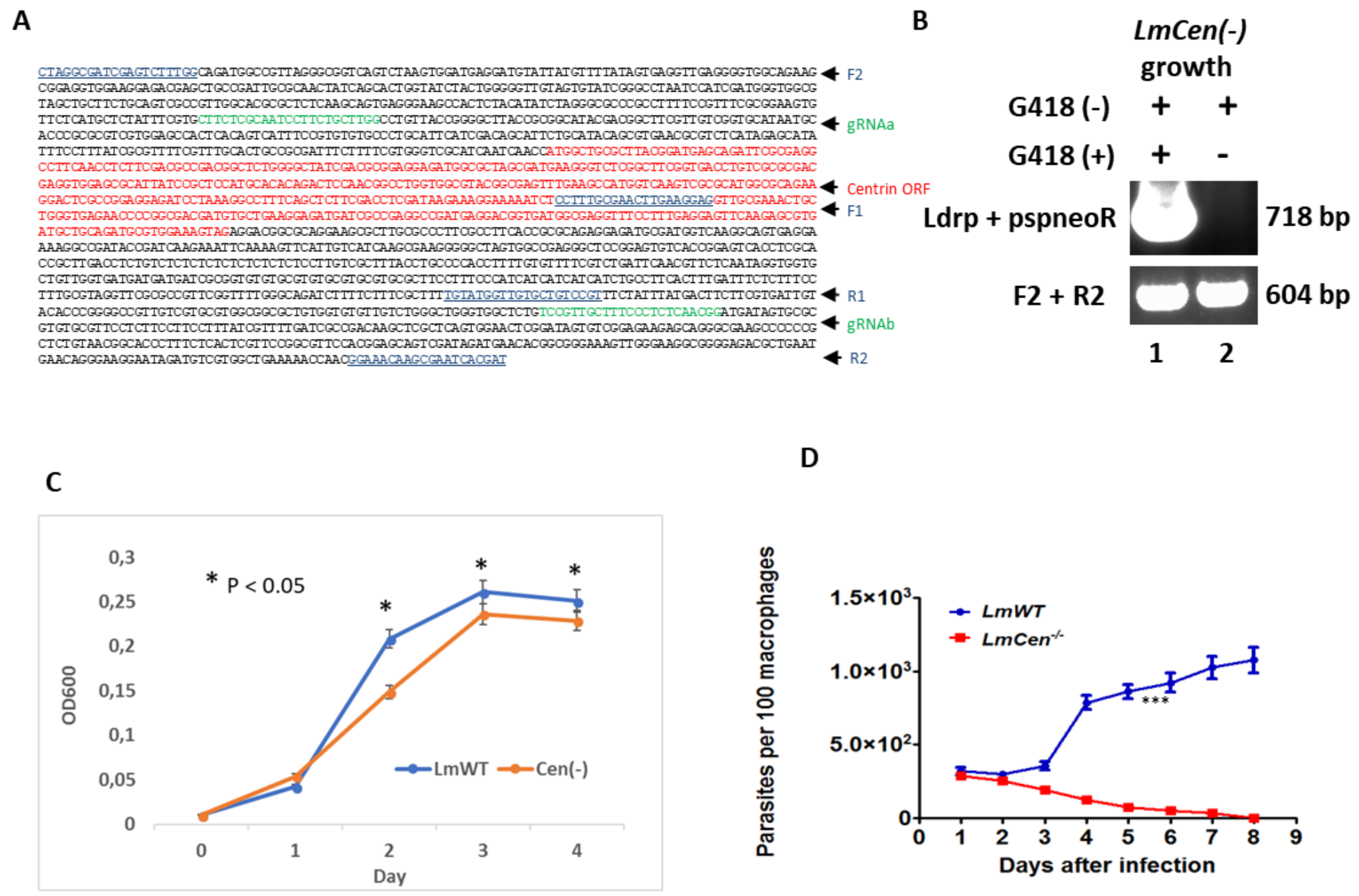


**Supplementary Figure 1**

**A.** *Leishmania major centrin* gene (LmjF.22.1410) and its flanking sequences. The CRISPR gRNAa and gRNAb targeting sites (Green) in the 5' and 3' *centrin* gene flanking sequences and PCR primers (Blue and underlined) used to confirm deletion of the *centrin* gene are indicated at the right.

**B.** Loss of antibiotic resistance CRISPR pLdCN plasmid in LmCen(-) cells after culture in G418 free medium for one month. To remove the antibiotic resistance CRISPR plasmid, these LmCen(-) cells were cultured in G418 free medium for several weeks before subject to cloning in a 96 well plate (free of G418), once the cell density in the 96 plate wells reached approximately 5 x $10^6$ per ml, 5 ul of the cell culture from each of the 96 plate wells was transferred to second 96 well plate with 100 ul medium per well containing 100 ug/ml G418. The genomic DNA extracted from LmCen-/- cells which were still able or not able to grow in G418 containing medium were subject to PCR analysis with the CRISPR plasmid specific primers (LdrP + pspneoR). No CRISPR plasmid specific PCR band was detected in the LmCen-/- cells which had lost the ability to grow in G418 containing culture medium, and the 604 bp F2+R2 band from the genome derived sequence could be detected in both G418 resistant and sensitive LmCen-/- cells.

**C.** LmCen-/- cells (Cen(-)) grow slower than wild-type *L. major* (LmWT) in promastigotes culture. L. major promastigotes were inoculated in a 96 well plate at the concentration of $10^6$ per ml, 120 ul per well and 5 wells per sample. The promastigotes growth was monitored by measuring the optical density in these wells at the wavelength of 600 nm (OD600) during the following 4 days. The data shown are the mean plus Standard Error of the Mean (SEM). Note, the cell density differences between LmWT and LmCen-/- cells at day 2,3 and 4 post inoculation are statistically significant ( $P < 0.05$). This is the representative data of three independent experiments.

D Human macrophages differentiated from monocytes were infected with stationary phase promastigote parasites from *LmWT* and *LmCen*$^{-/-}$ for six hours (10:1 parasite-to-macrophage ratio). The number of amastigotes in these cultures was determined over 8 days by microscopic observation of Diff-quik reagent stained slides. The data are expressed as the number of amastigotes per 100 macrophages. Error bars indicate the standard deviation (*** $p<0.001$).

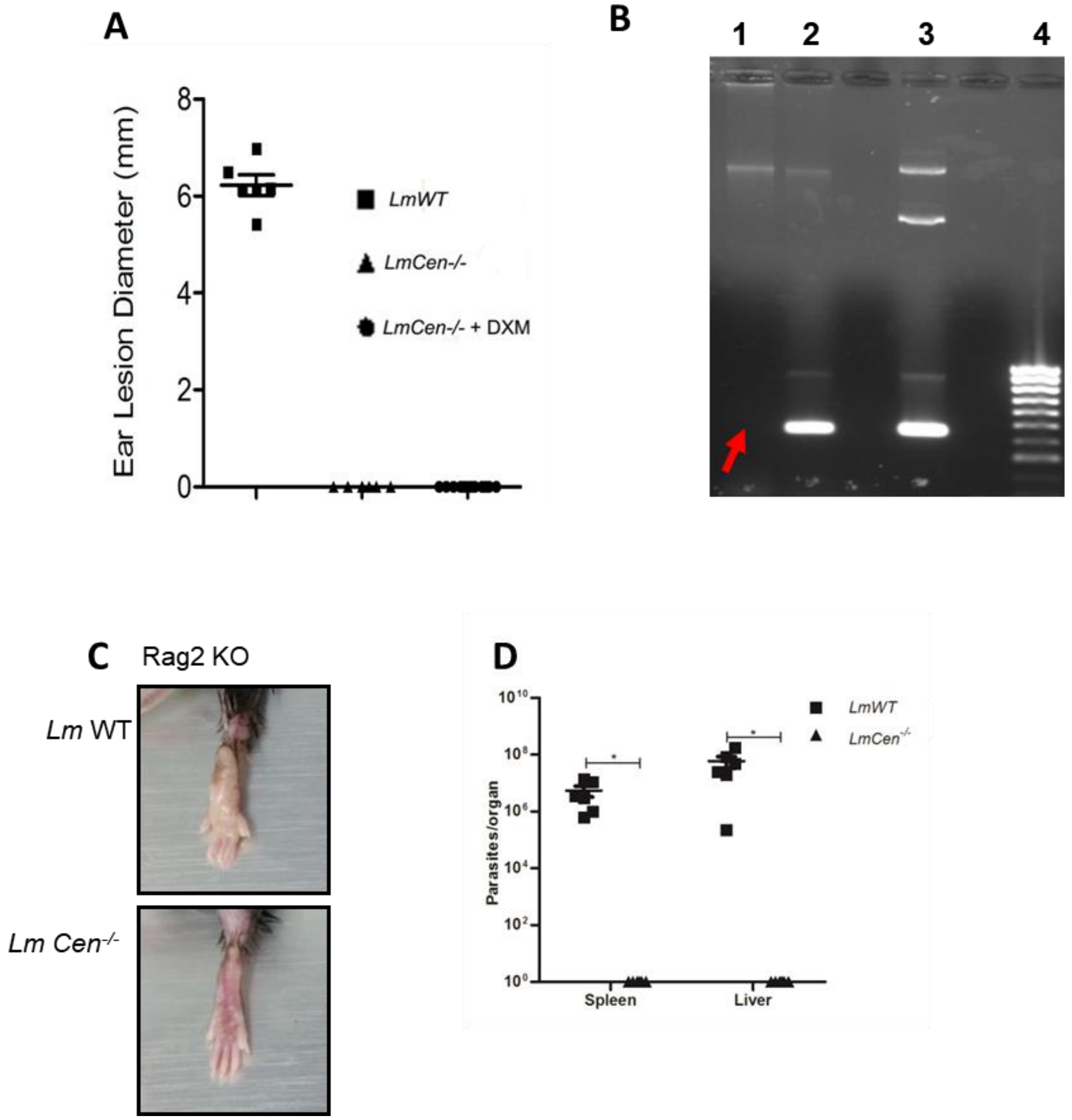


**Supplementary Figure 2**

**Safety and non-pathogenicity. A**. Ear lesion diameters was measured after 4 weeks of Dexamethasone (DXM) treatment (total 15 weeks post parasite infection) in *LmWT* and *LmCen*$^{-/-}$ (±DXM) immunized mice. Results are mean ± SEM, of 1 ear, 6-12 mice per group. **B.** 1% Agarose gel electrophoresis results for the characterization of *LmCen*$^{-/-}$ parasites isolated from *LmCen*$^{-/-}$ plus DXM treated group using *L. major centrin* gene specific primers. Lane-1, PCR results from the genomic DNA of parasites isolated from *LmCen*$^{-/-}$ plus DXM treated group, Lane-2, PCR results from the genomic DNA of parasites isolated from *LmWT* group, Lane-3, PCR results from the plasmid DNA containing *centrin* gene as a positive control . Lane 4 , 1 kb DNA ladder (Bioline). Red arrow indicates the absence of main product bands (*centrin* gene) of 450 bp in Lane-1. **C.** Representative photographs of footpad of Rag2 KO mice at 15 weeks post subcutaneous infection with $1 \times 10^7$ *LmCen*$^{-/-}$ or *LmWT*. D) Parasite burden in spleen and liver of Rag2 KO mice at 15 weeks post subcutaneous infection with $1 \times 10^7$ of *LmCen*$^{-/-}$ or *LmWT* into footpad.

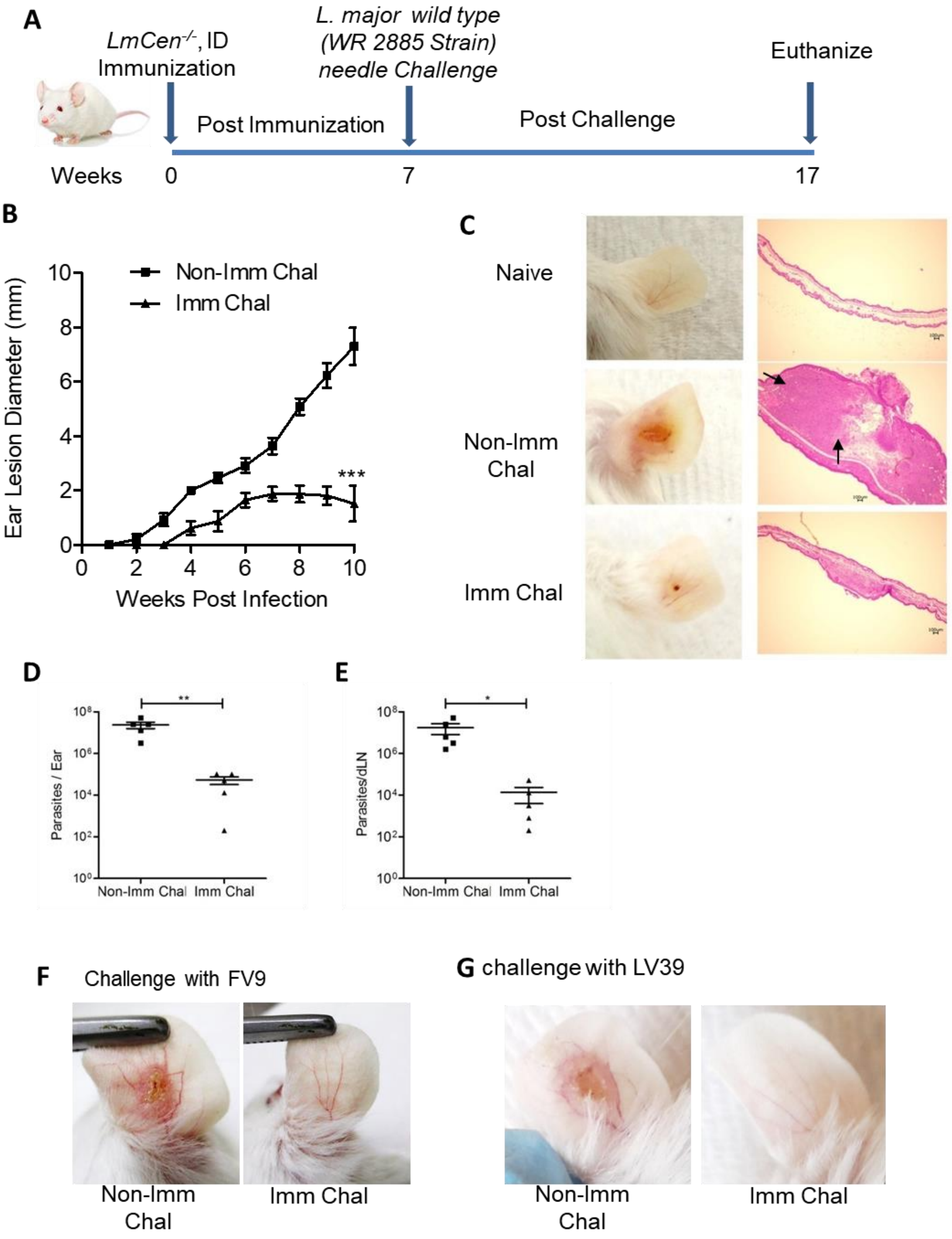


**Supplementary Figure 3**

**Protective efficacy of *LmCen*$^{-/-}$ parasites against virulent *L. major* needle challenge in BALB/c mice. A.** Schematic representation of the needle challenge procedure. BALB/c mice were immunized by intradermal injection in the left ear dermis with $1 \times 10^6$ stationary phase Centrin deleted *L. major* (*LmCen*$^{-/-}$) promastigotes. Seven weeks post-immunization, both immunized & age matched naïve animals were challenged with 750 metacyclic *L. major* wild type parasites (strain WR2885) in the right ear by intradermal injection. All the animals were euthanized after 10 weeks post challenge as shown in the figure. **B.** Ear lesion size were measured weekly for both *LmCen*$^{-/-}$ immunized (Imm Chal) and non-immunized (Non-Imm Chal) mice after intradermal challenge with *LmWT* parasites. Results are mean ± SEM, of 1 ear, 5 mice per group. **C.** Photographs of representative challenged ear of *LmCen*$^{-/-}$ immunized (Imm Chal) & non immunized (Non-Imm Chal) mice after 10 weeks post challenge. Immunized challenged mice displaying significantly reduced ear lesion size and inflammation. Arrow indicates inflammatory cells recruited area. Parasite load of each *LmCen*$^{-/-}$ immunized (Imm Chal) & non-Immunized (Non-Imm chal) mice, ear (**D.**) and dLN (**E.**). Parasite burden was determined by limiting dilution assay. Results are representative of one independent experiment with five mice per group (females, 5–8 weeks old). Statistical analysis was performed by unpaired one-tailed t-test ($*p < 0.04$; $**p < 0.009$,$***p<0.0005$). **F.** BALB/c mice were immunized intradermally into ear with $1 \times 10^7$ *LmCen*$^{-/-}$ and, at 6 weeks post immunization, mice were challenged intradermally into alternate ear with 5,000 stationary phage FV9 *L. major* promastigotes.  Representative photographs of ear lesion of immunized (n=5) and non-immunized mice (n=5) at 14 weeks post challenge infection. **G.** BALB/c mice were immunized (n=5) subcutaneously in the footpad with 2 X $10^8$ *LmCen*$^{-/-}$ parasites; six weeks post immunization challenged intradermally into the ear with 10,000 LV39 *L. major* parasites Representative photographs of ear lesion of immunized and non-immunized mice at 10 weeks post challenge infection. All experiments were performed one time.

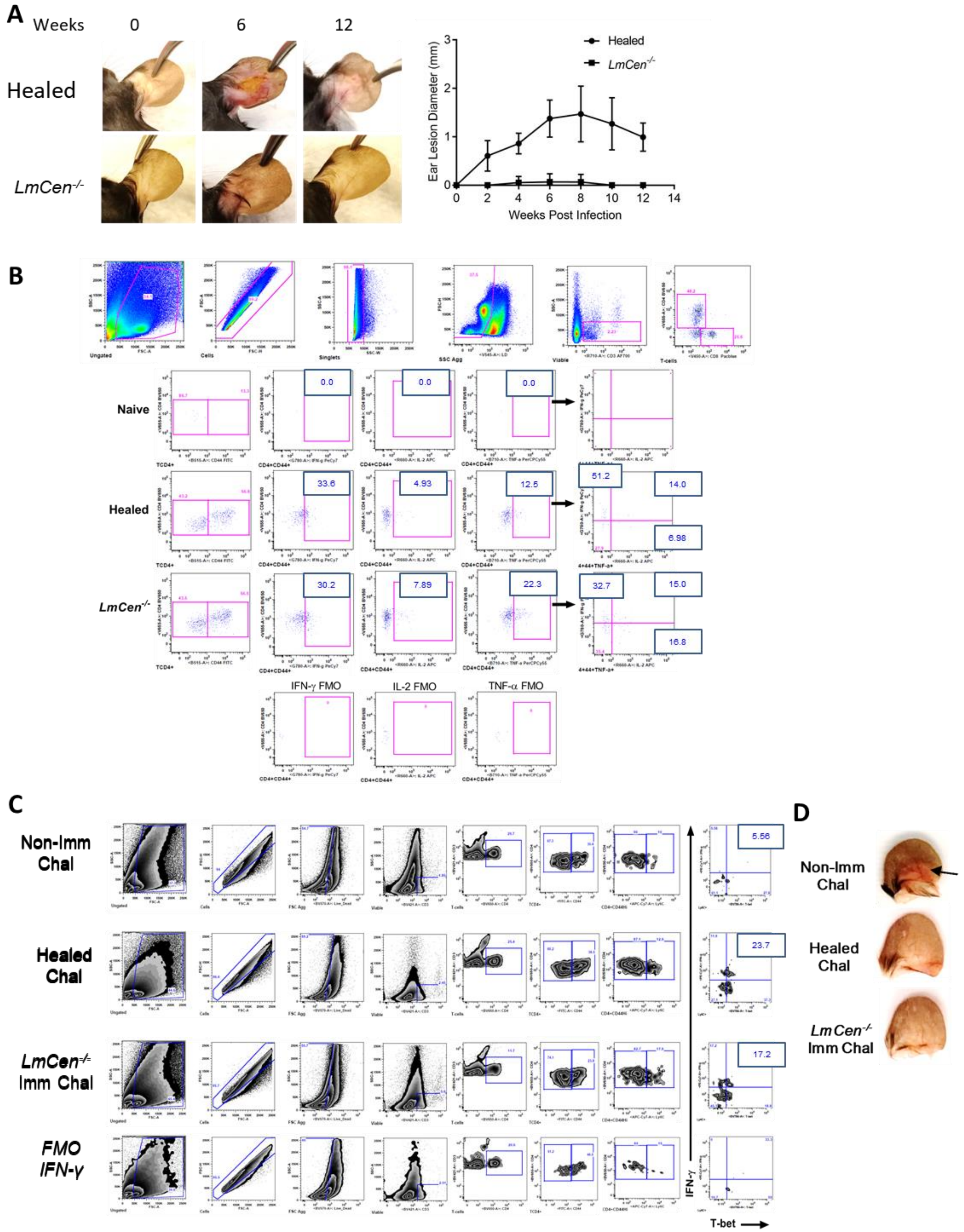


**Supplementary Figure 4**

***LmCen****$^{-/-}$* **immunization induces comparable host protective immune response with Healed from primary** ***LmWT*** **infection (leishmanization).** C57BL/6 mice were injected intradermally with $10^4$ metacyclic *LmWT* or $10^6$ total stationary phase *LmCen*$^{-/-}$ parasites and a comparative immune response between healed from primary *LmWT* infection (Leishmanized) and *LmCen*$^{-/-}$ immunized mice were determined following 8 weeks post-*LmCen*$^{-/-}$ immunization. To determine the 20h post challenge immune response, healed, *LmCen*$^{-/-}$ immunized as well as age matched naïve control mice were needle challenged with $10^5$ metacyclic *L. major* wild type (*LmWT*) parasites in the contralateral ear (at 12 weeks). **A.** Photograph of representative ear showing the course of lesion development and subsequent cure from primary *LmWT* infection in healed group of mice. Ear lesion size was measured weekly for both *LmCen*$^{-/-}$ immunized (*LmCen*$^{-/-}$) and healed from primary infected group (Healed) of mice after intradermal inoculation of parasites. Results are mean ± SEM, of 1 ear, 5 mice per group. **B.** The common gating strategies and multiparameter flow cytometry based analysis for single, double or triple cytokine secreting $CD4^+$ $CD44^+$ T cells after 20h *in-vitro* re-stimulation with freeze-thaw *L. major* antigen (*LmFTAg*) from pooled ear of naïve (no immunization and no challenged), healed and *LmCen* $^{-/-}$ immunized group of mice plus naive splenic APCs. Antigen experienced cells were gated and divided into six distinct subpopulations and the percentage of the various subpopulations were calculated. **C.** Common gating steps and representative zebra plots of early immune response in the ear of age matched non-immunized, healed and *LmCen*$^{-/-}$ immunized mice following needle challenge with wild type *L. major*-parasites. Naïve mice injected with PBS is control group. Twenty-hour post-challenge, ear-derived cells were analyzed and represented as the percentage of IFN-γ-producing $CD4^+CD44^{Hi}$T-bet$^+$Ly-6C$^+$ -T cells in response to 12-14 hours *in-vitro* re-stimulation with freeze-thaw *L. major* antigen (*Lm*Ag) plus naive splenic APCs. **D.** Representative photographs of ear lesion of age matched non immunized, healed and *LmCen*$^{-/-}$ immunized mice at 5 weeks post *LmWT* infected sand fly challenge. Only age matched non immunized group develop lesion (black arrow) at 5 weeks post challenge compared to healed and *LmCen*$^{-/-}$ immunized mice. Positivity for each antibody against intra cellular cytokines was determined using fluorescence minus one (FMO) controls.
